# Opposing immune and genetic forces shape oncogenic programs in synovial sarcoma

**DOI:** 10.1101/724302

**Authors:** Livnat Jerby-Arnon, Cyril Neftel, Marni E. Shore, Matthew J. McBride, Brian Haas, Benjamin Izar, Hannah R. Weissman, Angela Volorio, Gaylor Boulay, Luisa Cironi, Alyssa R. Richman, Liliane C. Broye, Joseph M. Gurski, Christina C. Luo, Ravindra Mylvaganam, Lan Nguyen, Shaolin Mei, Johannes c. Melms, Christophe Georgescu, Ofir Cohen, Jorge E. Buendia-Buendia, Michael S. Cuoco, Danny Labes, Daniel R. Zollinger, Joseph M. Beechem, G. Petur Nielsen, Ivan Chebib, Gregory Cote, Edwin Choy, Igor Letovanec, Stéphane Cherix, Nikhil Wagle, Peter K. Sorger, Alex B. Haynes, John T. Mullen, Ivan Stamenkovic, Miguel N. Rivera, Cigall Kadoch, Orit Rozenblatt-Rosen, Mario L. Suvà, Nicolò Riggi, Aviv Regev

**Affiliations:** Broad Institute of Harvard and MIT, Cambridge, MA, 02142, USA; Klarman Cell Observatory, Broad Institute of Harvard and MIT, Cambridge, MA 02142, USA; Department of Pathology and Center for Cancer Research, Massachusetts General Hospital and Harvard Medical School, Boston, MA, 02114, USA; Institute of Pathology, Faculty of Biology and Medicine, Centre Hospitalier Universitaire Vaudois, Lausanne, 1011, Switzerland; Department of Pediatric Oncology, Dana-Farber Cancer Institute and Harvard Medical School, 450 Brookline Avenue, Boston, MA, 02215, USA; Department of Medical Oncology, Dana-Farber Cancer Institute and Harvard Medical School, 450 Brookline Avenue, Boston, MA, 02215, USA; Massachusetts General Hospital Cancer Center, 55 Fruit Street, Boston, MA, 02114, USA; Laboratory for Systems Pharmacology, Harvard Medical School, Boston, MA, 02115, USA; Flow Cytometry Facility, Department of Biology and Medicine, University of Lausanne, Lausanne, 1011, Switzerland; NanoString Technologies Inc., 530 Fairview Avenue North, Seattle, WA 98109, USA; Department of Medicine, Division of Hematology and Oncology, Massachusetts General Hospital, Boston, MA, 02114, USA; Department of Orthopedics, Faculty of Biology and Medicine, Centre Hospitalier Universitaire Vaudois, Lausanne, 1011, Switzerland; Department of Surgery, Massachusetts General Hospital, Boston, MA, 02114, USA; Howard Hughes Medical Institute, Koch Institute for Integrative Cancer Research, Department of Biology, MIT, Cambridge, MA, 02139, USA

## Abstract

Synovial sarcoma is an aggressive mesenchymal neoplasm, driven by the SS18-SSX fusion, and characterized by immunogenic antigens expression and exceptionally low T cell infiltration levels. To study the cancer-immune interplay in this disease, we profiled 16,872 cells from 12 human synovial sarcoma tumors using single-cell RNA-sequencing (scRNA-Seq). Synovial sarcoma manifests antitumor immunity, high cellular plasticity and a core oncogenic program, which is predictive of low immune levels and poor clinical outcomes. Using genetic and pharmacological perturbations, we demonstrate that the program is controlled by the SS18-SSX driver and repressed by cytokines secreted by macrophages and T cells in the tumor microenvironment. Network modeling predicted that SS18-SSX promotes the program through HDAC1 and CDK6. Indeed, the combination of HDAC and CDK4/6 inhibitors represses the program, induces immunogenic cell states, and selectively targets synovial sarcoma cells. Our study demonstrates that immune evasion, cellular plasticity, and cell cycle are co-regulated and can be co-targeted in synovial sarcoma and potentially in other malignancies.

## INTRODUCTION

Therapeutic strategies harnessing the cytotoxic capacity of the adaptive immune response to target tumor cells have radically changed clinical practice, but response varies dramatically across patients and tumor types (1,2). Cancer types with defined genetics and exceptionally low T cell infiltration levels could help provide clues to some of the immune escape mechanisms underlying lack of response to immune therapies.

One such cancer type is synovial sarcoma (SyS) (3), a highly aggressive mesenchymal neoplasm that accounts for 10-20% of all soft-tissue sarcomas in young adults (4). SyS tumors homogenously express several immunogenic cancer-testis antigens (CTAs) (5–8), which are recognized by circulating T cells in the peripheral blood of SyS patients (5–7). Nonetheless, T cell infiltration remains exceptionally low in these tumors, suggestive of yet unidentified immune evasion mechanisms.

The cellular plasticity (4), stem-like features (9,10), and unique genetics of SyS may explain its exceptional ability to escape immune surveillance despite the expression of immunogenic antigens. SyS is invariably driven by the SS18-SSX oncoprotein – where the BAF subunit SS18 is fused to SSX1, SSX2 or, rarely, SSX4 (11). The BAF complex, the mammalian ortholog of SWI/SNF, is a major chromatin regulator (11), which has been previously shown to mediate resistance to immune checkpoint blockade in melanoma and renal cancer (12,13). SSX genes are a family of CTAs involved in transcriptional repression (14–17). The resulting SS18-SSX oncoprotein leads to massive dysregulation of the chromatin architecture and transcriptional regulation (11,18–20), generating a spectrum of malignant cellular morphologies (4), including epithelial-like malignant cells (in biphasic tumors), suggestive of pluripotential differentiation or mesenchymal to epithelial transitions.

Studies of human SyS to date have either relied on bulk tissue (21,22) or on established cellular models (11,18,19), masking important aspects of the tumor ecosystem. Moreover, given this cancer’s rarity, even concerted efforts, such as TCGA, profiled only limited numbers of tumors (21–23). Here, we leveraged single-cell RNA-Seq (scRNA-Seq), imaging, functional perturbations, and computational modeling, to study the cancer-immune interplay in SyS. We profiled 16,872 cells from 12 human SyS tumors by scRNA-seq and demonstrate that SyS tumors invariably include a subpopulation of cells expressing a novel core oncogenic program, associated with T cell exclusion. The core oncogenic program is predictive of poor prognosis and is repressed by the genetic inhibition of the SS18-SSX fusion, and by cytokines expressed by T cells and macrophages in the tumor microenvironment. HDAC1 and CDK6 are a key regulator and target of this aggressive cell program, respectively, and their combined inhibition synergistically represses it in SyS cells, triggering antigen presentation and cell autonomous immune responses. Collectively, our findings demonstrate a strong connection between SyS development and immune evasion, and strengthen the notion that de-differentiation, immune evasion, and cell cycle are co-regulated, such that cellular immunity can be targeted through modulation of cell cycle and epigenetic processes.

## RESULTS

### Cell type inference from expression and genetic features in scRNA-seq of SyS

To comprehensively interrogate the SyS ecosystem, we used full-length (24) and droplet-based (25) scRNA-Seq to profile 16,872 high quality malignant, immune, and stromal cells from 12 human SyS tumors (**Fig. 1A,B, Supp. Fig. 1A,B, Supp. Table 1, Methods**). We assigned cells to different cell types according to both genetic and transcriptional features (**Fig. 1B-G, Supp. Fig. 1, Methods**): (1) expression-based clustering and *post hoc* annotation of non-malignant clusters based on canonical cell type markers (**Fig. 1C, Supp. Fig. 1A, Supp. Table 2**); (2) detection of the *SS18-SSX* fusion transcripts (26) (**Fig. 1D**); (3) inference of copy number alterations (CNAs) from scRNA-Seq profiles (27) (**Fig. 1E**), which we validated in four tumors by bulk whole-exome sequencing (WES) (**Fig. 1G**); and (4) similarity of cells to bulk expression profiles of SyS tumors (**Methods**) (23) (**Fig. 1F**). The four approaches were highly congruent (**Supp. Fig. 1A**). For example, the fusion was detected in 58.6% of cells inferred as malignant by other analyses, but only in 0.89% of non-malignant cells. Notably, *SSX1/2* expression was also very specific to malignant cells, with a detection rate of 66.64% and 1.49% in the malignant and non-malignant cells, respectively (**Supp. Fig. 1A**, “SSX1/2 detection”). Similarly, CNAs were detected only in cells that were assigned as malignant by the other analyses (**Fig. 1E,G**), and the SyS similarity scores distinguished between malignant and non-malignant cells (as defined by the other methods) with 100% accuracy (**Fig. 1F, Supp. Fig. 1C**). Cells discordant across these criteria (< 0.05%) were excluded from all downstream analyses. Notably, in one of the tumors we identified an additional malignant-specific fusion between *MEOX2* and *AGMO* (**Supp. Fig. 1A**).

**Figure 1.**
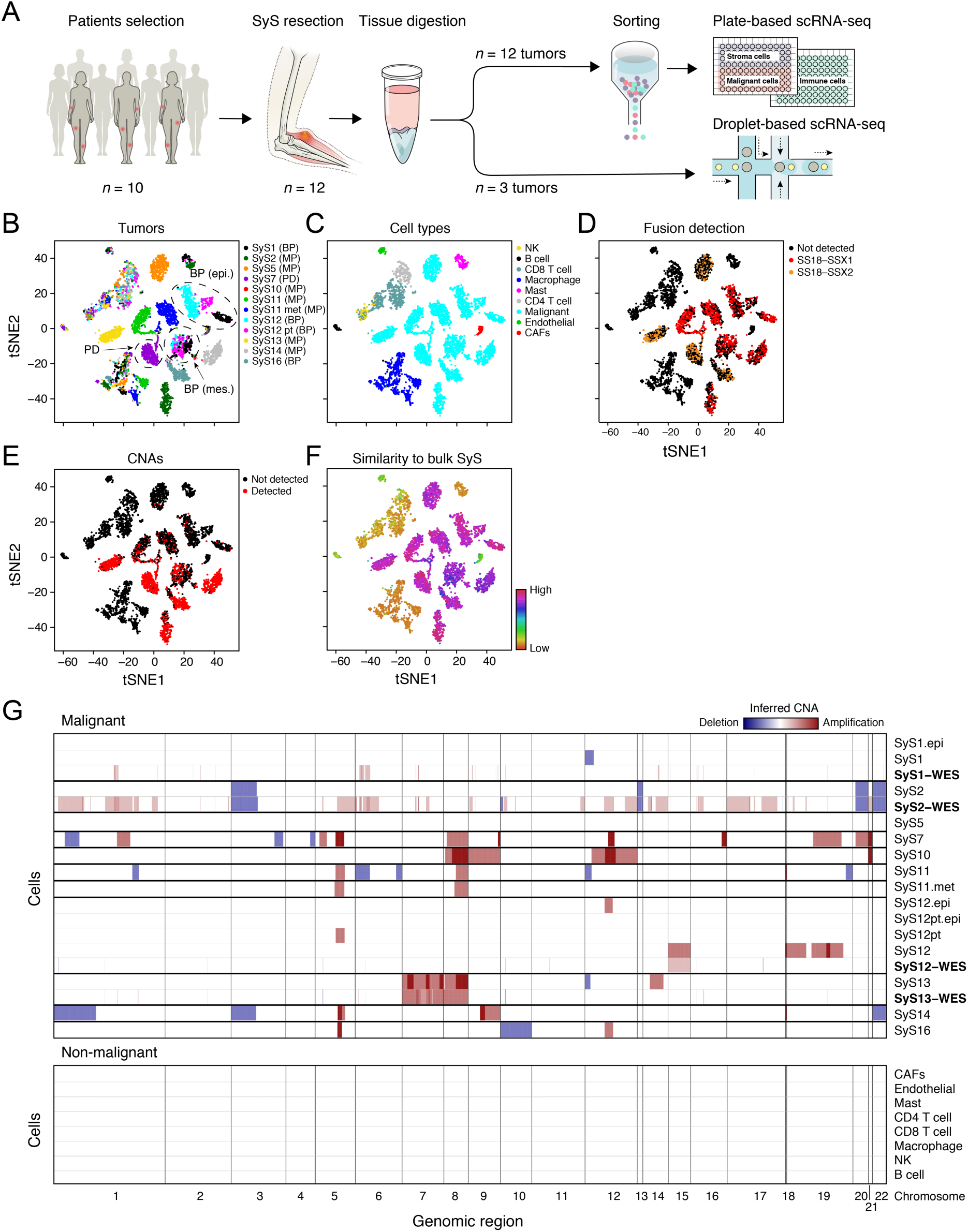
A single-cell map of the cellular ecosystem of synovial sarcoma tumors. **(a)** Study workflow. **(b-e)** Consistent assignment of cell identity. t-SNE plots of scRNA-Seq profiles (dots), colored by either **(b)** tumor sample, **(c)** inferred cell type, **(d)** SS18-SSX1/2 fusion detection, **(e)** CNA detection, and **(f)** differential similarity to SyS compared to other sarcomas (**Methods**). Dashed ovals (b): mesenchymal and epithelial malignant subpopulations of biphasic (BP) tumors or poorly differentiated (PD) tumor. **(g)** Inferred large-scale CNAs distinguish malignant (top) from non-malignant (bottom) cells, and are concordant with WES data (bold). The CNAs (red: amplifications, blue: deletions) are shown along the chromosomes (x axis) for each cell (y axis).

We assigned the cells to nine subsets (**Fig. 1C**): malignant cells, non-malignant endothelial cells, Cancer Associated Fibroblasts (CAFs), CD8 and CD4 T cells, B cells, Natural Killer (NK) cells, macrophages, and mastocytes, and generated signatures for each subset (**Supp. Table 2, Supp. Fig. 1D**). Malignant cells primarily grouped by their tumor of origin, while their non-malignant counterparts (immune and stroma) grouped primarily by cell type (**Fig. 1B,C**), as we have observed in other tumor types (28–30). The malignant cells of each of the biphasic (BP) tumors (SyS1 and SyS12) formed two distinct subsets – epithelial and mesenchymal – which clustered together with malignant cells of the other biphasic tumor (**Fig. 1B,C**, black, cyan and magenta dots, **Methods**). We next focused on characterizing the states of immune cells in SyS.

### Evidence of antitumor immune activity despite low immune infiltration

The lack of effective antitumor immunity in SyS may results from: either the inactivity of immune cells, limiting their recognition of or response to SyS malignant cells, or hampered immune cell infiltration and recruitment into the tumor parenchyma. To test the first possibility, we examined CD8 T cell states (**Fig. 2A, Supp. Table 3A**), and found clear hallmarks of antitumor immunity and recognition. T cell subsets span naïve, cytotoxic, exhausted, and regulatory T cells (**Fig. 2B; Methods**), with evidence of expansion based on TCR reconstruction (31) (showing 57 clones, all patient-specific, with shared clones between matched samples from the same patient). While cytotoxic and exhaustion markers were generally co-expressed in T cells (**Fig. 2B**, consistent with previous reports (29)), clonally expanded T cells had unique transcriptional features (**Methods, Supp. Table 3A**), suggestive of an effector-like non-exhausted state (**Fig. 2B**, P < 6.6*10^-12^, mixed-effects). These expanded T cells might respond to SyS-specific CTAs, which were specifically expressed in large fractions of the malignant cell populations (**Supp. Fig. 2A**). Moreover, CD8 T cells in SyS have features suggesting they are even more active than those in melanoma tumors, where anti-tumor immunity is relatively pronounced. First, compared to CD8 T cells from melanoma (32), CD8 T cells in SyS tumors overexpressed a program characterizing T cells in tumors that were responsive to immune checkpoint blockade (33) (**Fig. 2C** bottom, P = 1.22*10^-10^, mixed-effects). In addition, compared to melanoma CD8 T cells, the SyS CD8 T cells also overexpressed effector and cytotoxic gene modules (34,35) (*e.g., GZMB, CX3CR1*, P = 6.36*10^-9^, mixed-effects), and repressed exhaustion markers (P = 6.36*10^-3^, mixed-effects), including *LAYN* (34), and multiple checkpoint genes (*CTLA4, HAVCR2, LAG3, PDCD1*, and *TIGIT*; P = 7.69*10^-7^, mixed-effects, **Fig. 2C**, top).

**Figure 2.**
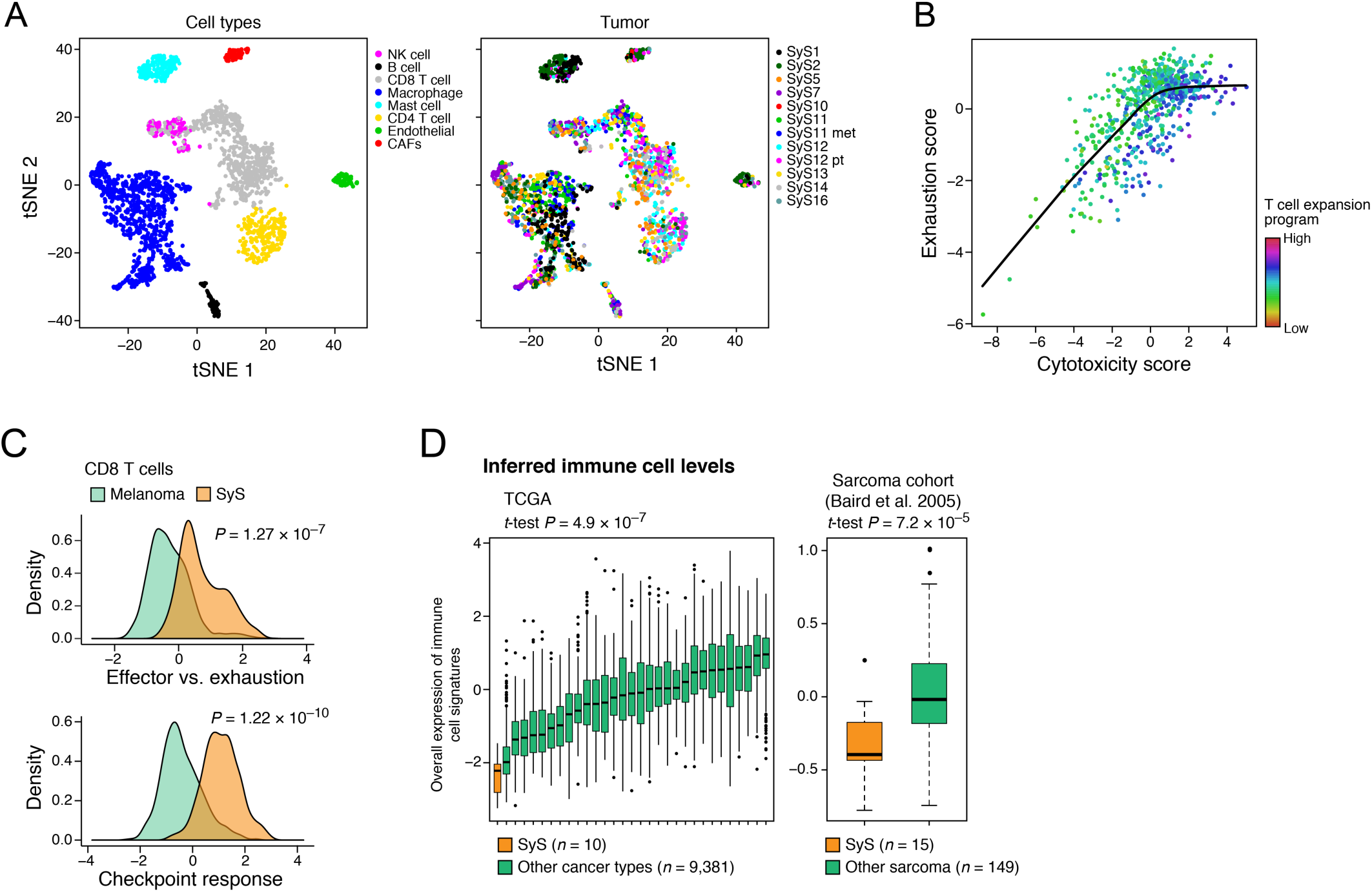
SyS tumors manifest antitumor immunity with limited immune infiltration. **(a)** Immune and stroma cells in SyS tumors. t-SNE of immune and stroma cell profiles (dots), colored by inferred cell type (left) or sample (right). **(b)** The CD8 T cell expansion program is associated with particularly high cytotoxicity and lower than expected exhaustion. The cytotoxicity (x axis) and exhaustion (y axis) scores of SyS CD8 T cells, colored by the score of the T cell expansion program (**Methods**). **(c)** CD8 T cells in SyS (orange) have higher effector programs than in melanoma (green). Distribution of effector *vs*. exhaustion scores (*x* axis, top, **Methods**) or an immune checkpoint blockade responsiveness program (33) (*x* axis, bottom, **Methods**) in CD8 T cells from each cancer type. **(d)** SyS tumors manifest a particularly cold phenotype. Overall Expression of the immune cell signatures (*y* axis, **Methods**) in SyS tumors (orange) and other cancer types (left panel) or other sarcomas (right panel).

Other immune cells in the tumor microenvironment also showed features of antitumor immunity. Macrophages span M1-like and M2-like states, suggestive of both pro- and anti-inflammatory properties, respectively (**Supp. Fig. 2B-D**; **Methods, Supp. Table 3B**), and expressed relatively high levels of TNF (P = 1.13*10^-7^, mixed-effects, >4 fold more compared to melanoma macrophages). However, mastocytes show regulatory features, with 39% of them expressing PD-L1 (as opposed to only 2% PD-L1 expressing malignant cells).

We next examined the alternative hypothesis that T cell abundance might be a limiting factor in SyS, despite these favorable T cell states. We compared SyS to 30 other cancer and sarcoma types. SyS tumors showed extremely low levels of immune cells, which cannot be explained by variation in the mutational load (**Fig. 2D**; P = 2.58*10^-11^, mixed effects when conditioning on the tumor mutational load), and despite the malignant-cell specific expression of immunogenic CTAs (**Supp. Fig. 2A**). In addition, unlike melanoma (**Supp. Fig. 2E**, left), T cell levels were not correlated with prognosis in SyS (**Supp. Fig. 2E**, right), indicating that they may not cross the critical threshold to impact clinical outcomes. Only mastocytes had a moderate positive association with improved prognosis (P = 0.012, Cox regression). These findings suggest that the lack of proper immune cell recruitment and infiltration is a key immune evasion mechanism in SyS, potentially mediated by the SyS cells.

### Cellular plasticity and a core oncogenic program characterize synovial sarcoma cells

To identify malignant cell functions that may impact immune cell infiltration, we characterized the cellular programs in SyS malignant cells. We identified three co-regulated gene modules, which repeatedly appeared across multiple tumors in our cohort (**Fig. 3A-D, Supp. Table 4, Methods**). The first two modules reflected mesenchymal and epithelial cell states (**Fig. 3B, Supp. Fig. 3A**). These differentiation programs included canonical mesenchymal (*ZEB1, ZEB2, PDGFRA* and *SNAI2*) or epithelial (*MUC1* and *EPCAM*) markers (36,37) (P < 1.55*10^-10^, hypergeometric test), and demonstrated that epithelial cells had a marked increase in antigen presentation and interferon (IFN) *γ* responses (P < 8.49*10^-6^, hypergeometric test).

**Figure 3.**
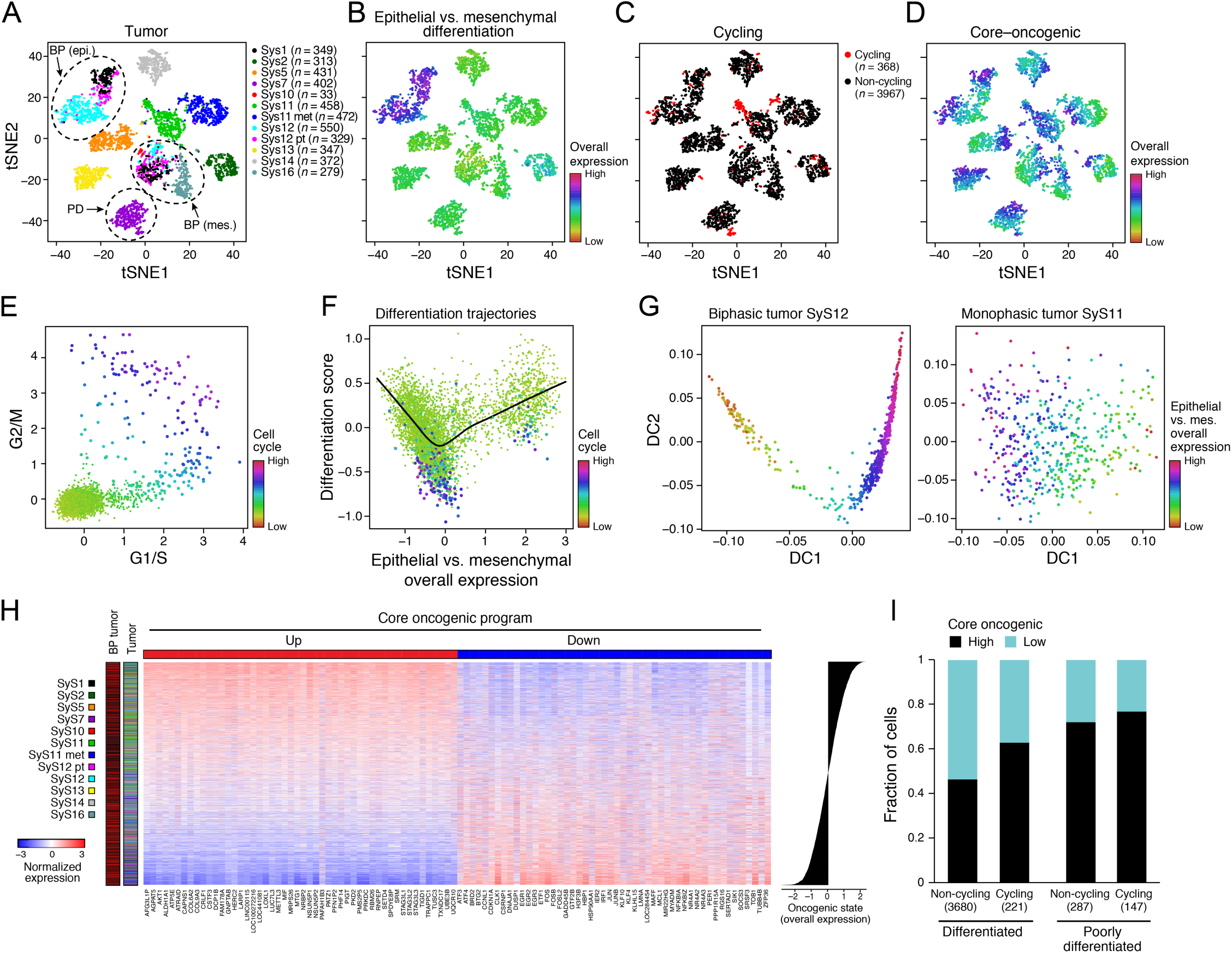
Cellular plasticity and a core oncogenic program characterize synovial sarcoma cells. **(a-d)** De-differentiation, cell cycle, and the core oncogenic programs across malignant cells. t-SNE plots of malignant cell profiles (dots), colored by: **(a)** sample, **(b)** Overall Expression of the epithelial *vs*. mesenchymal differentiation program, **(c)** cell cycle status, or **(d)** Overall Expression of the core oncogenic program. Dashed ovals (a): mesenchymal and epithelial malignant subpopulations of biphasic (BP) tumors or poorly differentiated (PD) tumor. **(e,f)** Association between cell cycle and poor differentiation. **(e)** G1/S (*x* axis) and G2/M (*y* axis) phase signature scores for each cell. **(f)** Epithelial and mesenchymal-like differentiation. Scatter plots of the malignant cells’ (dots) scores for the epithelial *vs*. mesenchymal program (*x* axis) and for overall differentiation (y axis). Color: expression of cell cycle program (see also **Supp. Fig. 3B,C**). **(g)** Distinct differentiation pattern in biphasic tumors. Single cell profiles dots arranged by the first two diffusion-map components (DCs) for representative examples of a biphasic (SyS12, left) and monophasic (SyS11, right) tumors, and colored by the Overall Expression of the epithelial *vs*. mesenchymal programs (colorbar). **(h)** Core oncogenic program genes. Normalized expression (centered TPM values, colorbar) of the top 100 genes in the core oncogenic program (columns) across the malignant cells (rows), sorted according to the Overall Expression of the program (bar plot, right). Leftmost color bars: biphasic tumor and sample ID. **(i)** The program is expressed in a higher proportion of cycling and poorly differentiated cells. Fraction of malignant cells (*y* axis) with a high (above median, black) and low (below median, blue) Overall Expression of the core oncogenic program, in cells stratified by cycling and differentiation status (*x* axis).

Among mesenchymal cells with a relatively low Overall Expression (**Methods**) of the mesenchymal program, one subset also expressed epithelial markers, reminiscent of transitioning to/from an epithelial state, while another underexpressed both programs, reminiscent of a poorly differentiated state. These poorly differentiated cells were highly enriched with cycling cells (P = 2.44*10^-60^, mixed effects), indicating that they might function as the tumor progenitors, fueling tumor growth (**Fig. 3E,F, Supp. Fig. 3B,C**). Diffusion map analysis of the cells based on these two programs highlighted putative differentiation trajectories, and found structured differentiation patterns only in the biphasic tumors (**Fig. 3G, Methods**). RNA velocity (38) demonstrated that epithelial to mesenchymal transitions may also take place (**Supp. Fig. 3D**), suggestive of cellular plasticity. Further supporting this hypothesis, the post-treatment sample of patient SyS12 includes a new subpopulation of mesenchymal cells, which was absent from the pre-treatment sample, and resembles the epithelial cells in terms of its CNAs (**Supp. Fig. 3E**).

The third module highlighted a new program present in a subset of cells in each tumor (25.2-84.7% per tumor, **Fig. 3D,H, Supp. Fig. 4**), which we named the core oncogenic program. The program is characterized by expression of genes from respiratory carbon metabolism (oxidative phosphorylation, citric acid cycle, and carbohydrate/protein metabolism, P < 1*10^-8^, hypergeometric test, **Supp. Table 4**), and repression of genes involved in TNF signaling, apoptosis, p53 signaling, and hypoxia processes (P < 1*10^-10^, hypergeometric test, **Supp. Table 4**), including known tumor suppressors, such as p21 (*CDKN1A*) and *KLF4*. The program was expressed in a higher proportion of cycling and poorly differentiated cells (P < 2.94*10^-4^, mixed-effects, **Fig. 3I**).

To test the clinical value of these transcriptional programs, we reanalyzed two independent bulk gene expression cohorts (21,22). Both dedifferentiation (**Methods**) and the core oncogenic program were substantially more pronounced in the more aggressive poorly differentiated SyS tumors (P < 2.76*10^-4^, one-sided t-test, **Fig. 4A, Methods**), and were associated with increased risk of metastatic disease (P < 1.36*10^-3^, Cox regression, **Fig. 4B**).

**Figure 4.**
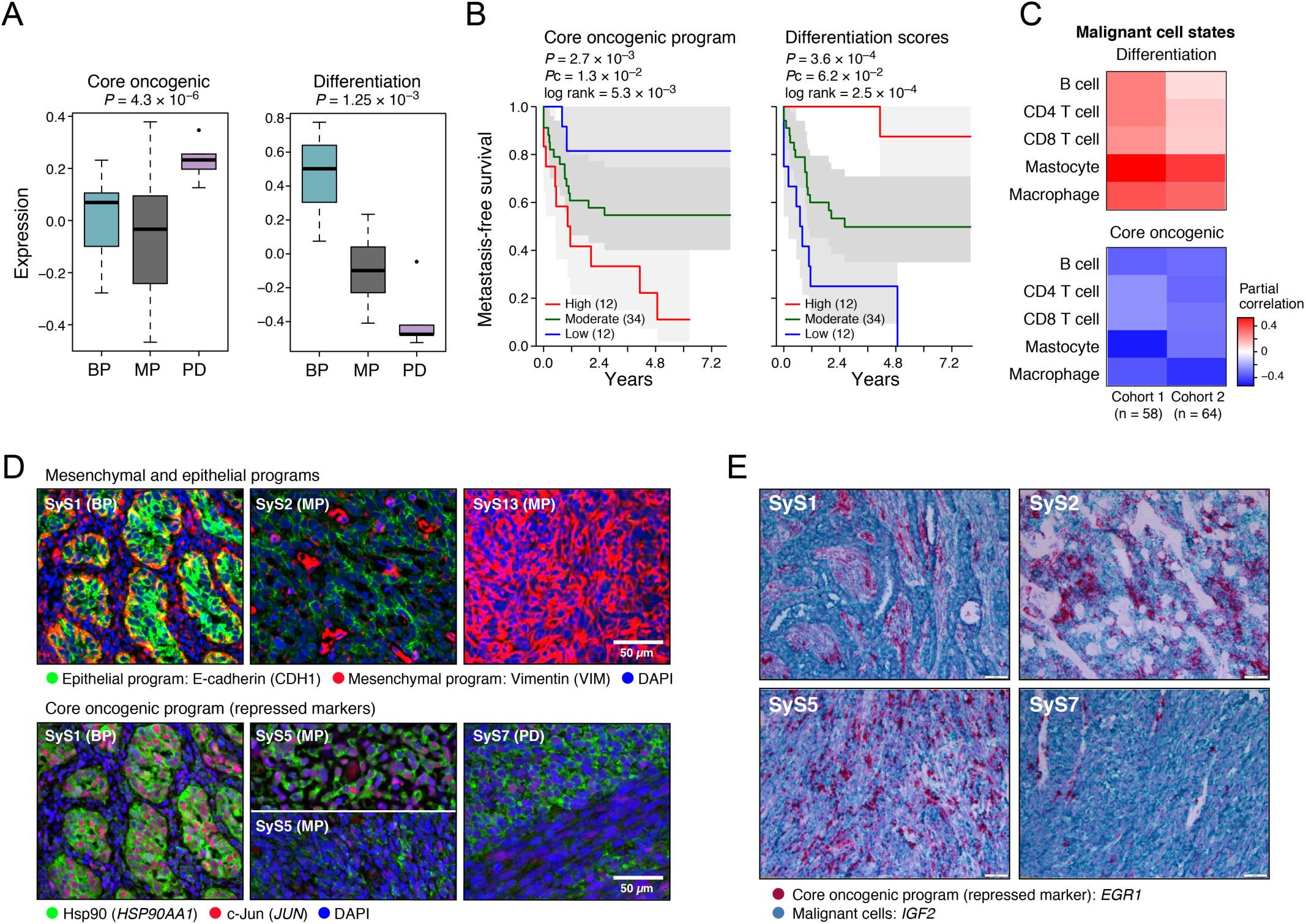
The core oncogenic program and de-differentiation co-vary within and across tumors and are associated with aggressive and cold tumors. **(a)** The core-oncogenic program and de-differentiation mark the aggressive poorly differentiated (PD) subtype. Overall expression of the core oncogenic or differentiation (both mesenchymal and epithelial) programs scores (y axis) across 34 SyS tumors (21), stratified as biphasic (BP), monophasic (MP), or poorly differentiated (PD) (x axis). Middle line: median; box edges: 25_th_ and 75_th_ percentiles, whiskers: most extreme points that do not exceed ±IQR*1.5; further outliers are marked individually. **(b)** The core oncogenic program and differentiation scores (overall expression of both differentiation programs) are predictive of metastatic disease in an independent cohort of 58 SyS patients (22). Kaplan-Meier (KM) curves of metastasis free survival (*x* axis, years), when stratifying the patients by high (top 25%), low (bottom 25%), or intermediate (remainder) expression of the respective program. *P*: COX regression p-value; *Pc:* COX regression p-value when controlling for fusion type and patient age group. **(c)** Inferred level of immune cell types is associated with the malignant programs in bulk SyS tumors, when controlling for tumor purity. Partial correlation (colorbar) between the inferred level of each immune subset (rows) and the core oncogenic and differentiation levels (columns). **(d-e)** *In situ* validation of programs. Detection of core oncogenic (Hsp90, c-Jun and *EGR1*), epithelial (E-cadherin) and mesenchymal (Vimentin) markers, using immunofluorescence (t-CyCIF) **(d)** and in situ hybridization (ISH) **(e)**.

### The core oncogenic program is associated with the cold phenotype and spatial niche

Next we turned to explore the connection between the malignant cells’ state and the tumor microenvironment and composition. Using our single-cell immune signatures we first estimated the composition of bulk SyS tumors in two published cohorts (18,22) and stratified them into “hot” or “cold”, based on their relative inferred proportions of immune cells (**Methods**). “Hot” tumors showed the repression of the core oncogenic program and had significantly higher differentiation scores (P < 5.34*10^-3^, *r* = −0.44 and 0.48, respectively, partial Pearson correlation, conditioning on inferred tumor purity, **Methods**; **Fig. 4C**).

Interestingly, the core oncogenic program shows some degree of similarity to a transcriptional signature we recently associated with T cell exclusion in melanoma (32) (P < 7.16*10^-10^, hypergeometric test), although most genes in the program (∼92%) were distinct from melanoma. Among the overlapping genes we find the induction of the CTA *MAGEA4*, the BAF complex unit *SMARCA4*, and repression of apoptosis and p53 signaling (e.g., *ATF3, JUN, KLF4*, and *SAT1*). The melanoma T cell exclusion signature and the synovial sarcoma mesenchymal state also overlapped (P = 6.33*10^-8^, hypergeometric test), for example, in the induction of *SNAI2* and repression of 23 epithelial genes, including *CDH1*. Nevertheless, the programs were largely distinct, likely given the different tissues, microenvironments, cell of origin and genetic drivers.

The association between the core oncogenic program and T cell exclusion is observed *in situ* in the SyS samples from our single-cell cohort. We measured *in situ* expression of 12 proteins across 4,310,120 cells in 9 samples using multiplexed immunofluorescence (t-CyCIF) (39) (**Fig. 4D,E**; **Methods**), and profiled the *in situ* expression of 1,412 genes in 24 spatially distinct areas in two samples using the GeoMx high plex RNA Assay (early version for Next-Generation Sequencing; **Methods**). Both approaches showed that CD45^+^ immune cells were exceptionally low in SyS (<0.4%, compared to >8.7% in melanoma samples (32)). Moreover, the malignant cells in the more immune infiltrated areas show a marked decrease in the core oncogenic program (*r* = −0.53, P = 6.9*10^-3^, Pearson correlation, and P < 1*10^-10^, mixed effects; **Methods**). This suggests that the status of the malignant cells and the composition of the tumor microenvironment might be interconnected in SyS.

### SS18-SSX sustains the core oncogenic program and blocks differentiation

To decouple the intrinsic and extrinsic factor determining the malignant cell states in SyS we first tested whether the core oncogenic and other programs were co-regulated by the genetic fusion driving SyS. We depleted SS18-SSX in two SyS cell lines (SYO1 and Aska) using shRNA, and profiled 12,263 cells by scRNA-Seq. The fusion knock-down (KD) led to extensive and highly consistent transcriptional alterations in both cell lines (**Fig. 5A, Supp. Fig. 5A, Supp. Table 5**): it substantially repressed the core oncogenic program and cell cycle genes (P < 8.05*10^-107^, t-test, **Fig. 5A-C**), while inducing mesenchymal differentiation programs and markers, including *ZEB1* and *VIM* (P < 1*10^-50^, t-test and likelihood-ratio test **Fig. 5A,B**,). The KD impact on the core oncogenic and differentiation programs was decoupled from the repression of cellular proliferation (**Fig. 5B**), such that the impact on the core oncogenic and differentiation programs was observed even when controlling for the cycling status of the cells, and when considering only cycling or non-cycling cells (P < 1.54*10^-13^, t-test, **Fig. 5B, Methods**). Thus, the fusion’s impact on cell cycle may be secondary or downstream to its impact on the core oncogenic program. In addition the fusion KD led to an induction of antigen presentation and cell autonomous immune responses, such as TNF and IFN signaling (P < 1*10^-30^, mixed-effects, **Supp. Fig. 5A**).

**Figure 5.**
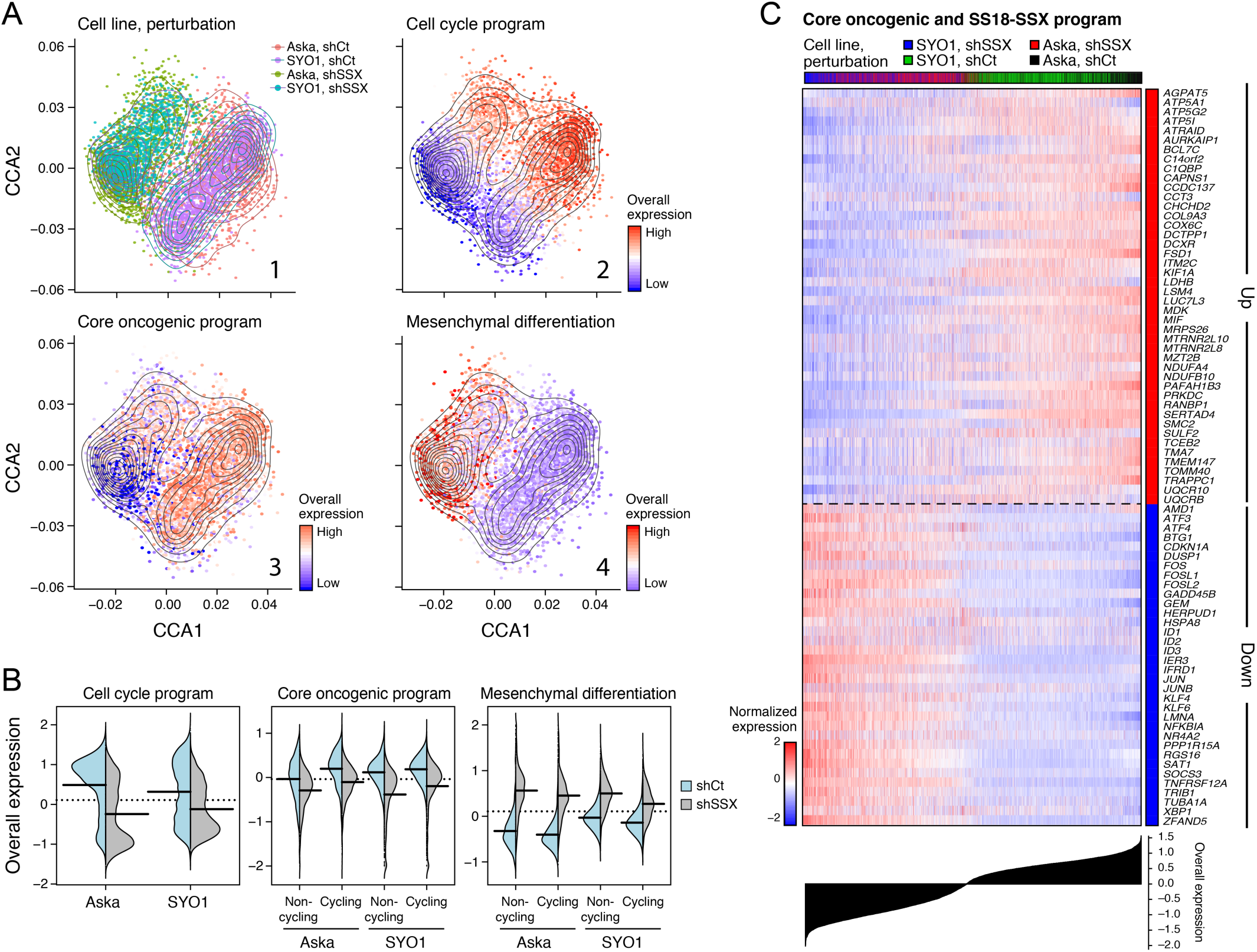
The genetic driver promotes the core oncogenic program in malignant SyS cells. **(a-c)** SS18-SSX sustains the core oncogenic, cell cycle, and dedifferentiation programs. **(a)** scRNA-Seq following KD of SS18-SSX. Co-embedding (using PCA and canonical correlation analyses (48), **Methods**) of Aska and SYO1 cell profiles (dots), colored by: (1) cell line and perturbation; or the Overall Expression (colorbar) of the (2) cell cycle, (3) core oncogenic, or (4) mesenchymal differentiation (36,37) programs. **(b)** SS18-SSX KD represses the core oncogenic program and induces the mesenchymal differentiation program irrespective of its repression of the cell cycle program. Distribution of Overall Expression scores (y axis) for each program in control (blue) and shSSX (grey) cells, for each cell line, where core oncogenic and mesenchymal program scores are shown separately for cycling and non-cycling cells. **(c)** Overlap of SS18-SSX and core oncogenic programs. Expression (centered TPM) of genes (rows) shared between the fusion and core oncogenic programs across the Aska and SYO1 cells (columns), with a control (shCt) or SSX (shSSX) shRNA. Cells are ordered by the Overall Expression of the SS18-SSX program (bottom plot) and labeled by type and condition (Color bar, top).

Using these SS18-SSX KD experiments we defined an SS18-SSX program, which we then stratified to direct and indirect fusion targets based on available SS18-SSX ChIP-Seq profiles (*13, 28*) (**Methods; Supp. Fig. 5B,C, Supp. Table 5A**). According to the SS18-SSX program, the fusion directly dysregulates developmental programs and promotes the core oncogenic program (P < 2.51*10^-5^, hypergeometric test, **Methods, Supp. Fig. 5B, Supp. Table 5**), while its impact on cell cycle genes is mostly indirect (P < 1.2*10^-9^, hypergeometric test, **Supp. Table 5, Supp. Fig. 5B**) and mediated by cyclin D2 (*CCND2*) and *CDK6* – the only cell cycle genes that are members of the direct SS18-SSX program. Taken together, our findings support a model in which SS18-SSX directly promotes the core oncogenic program, blocks differentiations, drives cell cycle progression, and represses features necessary for immune recognition and recruitment.

### TNF and IFN*γ* synergistically repress the core oncogenic and SS18-SSX programs

The association between the core oncogenic program and the cold phenotype suggest that the program promotes T cell exclusion in SyS. Another (non-mutually exclusive) hypothesis is that, despite their low numbers, the immune cells in the tumor microenvironment may nonetheless impact the state of the malignant cells, for example, through the secretion of different molecules and cytokines. To test this, we implemented a mixed-effects inference approach that uses scRNA-Seq data to find associations between the expression of secreted molecules and ligands in immune cells and the state of the malignant cells (**Methods**).

According to this analysis, the expression of IFN*γ* and TNF specifically from CD8 T cells and macrophages, respectively (**Fig. 6A**), was strongly associated with the repression of the core oncogenic program in the malignant cells (P < 9.4*10^-39^, mixed-effects). We further stratified the core oncogenic program to its predicted TNF/IFN*γ*-dependent and -independent components, by the association of each gene’s expression in the malignant cells with the TNF and IFN*γ* expression levels in the corresponding macrophages and CD8 T cells, respectively (**Methods, Supp. Table 6A**).

**Figure 6.**
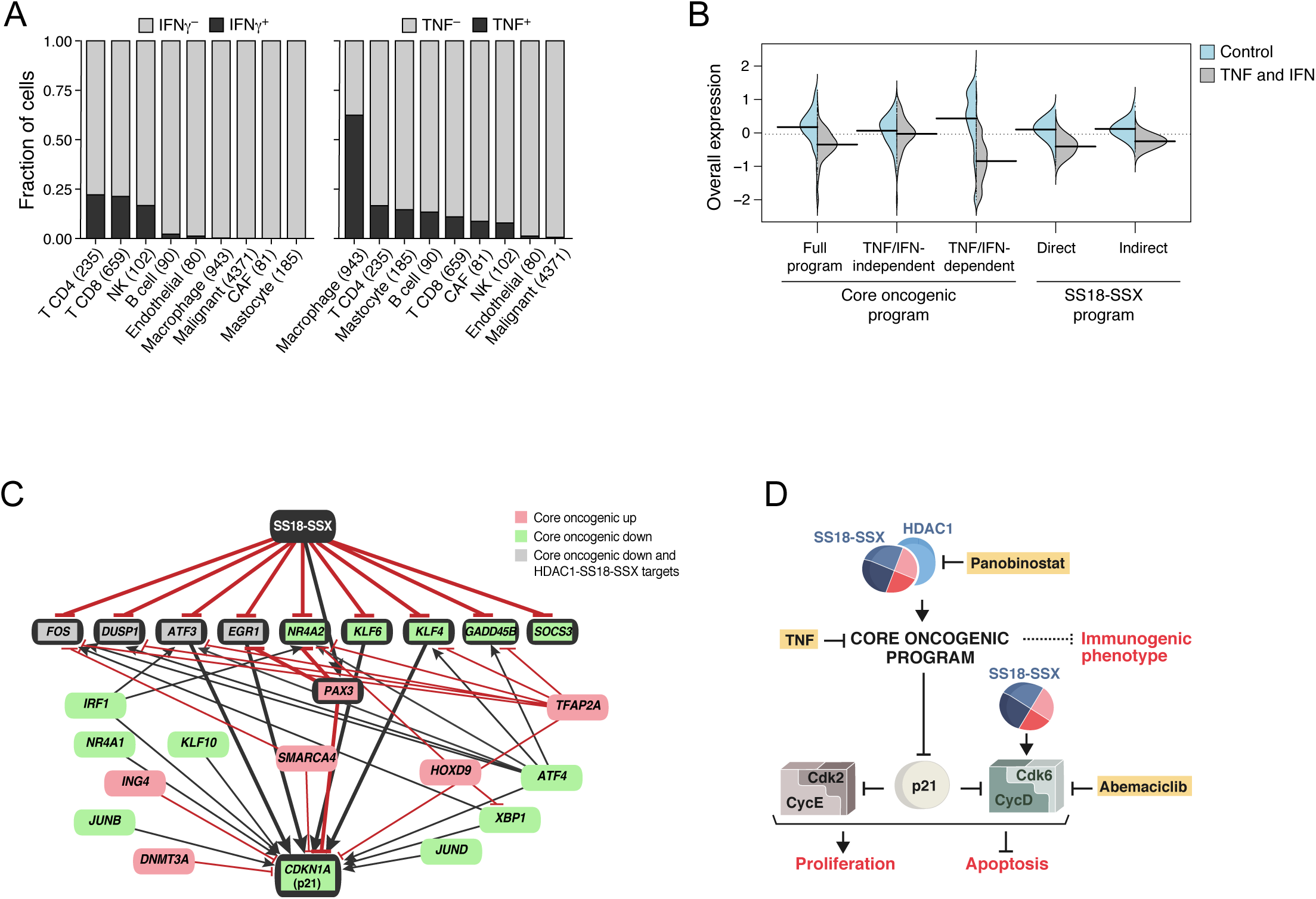
The immune cells and the genetic driver form two opposing forces in shaping SyS cell states. **(a,b)** TNF and IFN*γ* are expressed by immune cells in the tumor microenvironment and can repress the core oncogenic program. **(a)** TNF and IFN*γ* are detected primarily in macrophages and T cells, respectively. Fraction of cell (y axis) of each subset in the tumor (x axis) that express (black) IFN*γ* (left) or TNF (right) by scRNA-seq. **(b)** TNF and IFN*γ* repress the core oncogenic and SS18-SSX programs (see also **Supp. Fig. 5D**). Distribution of Overall Expression score (y axis) of the core oncogenic (also stratified to its predicted and TNF/IFN*γ*-dependent and -independent components) and SS18-SSX programs (x axis) in control (blue) and TNF + IFN*γ* treated cells. **(c)** Gene regulatory model of control of the core oncogenic program by SS18-SSX. Red/green: genes that are induced/repressed in the core oncogenic program. Grey: genes that are repressed in the core oncogenic program and directly repressed by HDAC1-SS18-SSX (20). Red blunt arrows: repression; black pointy arrows: activation. Thick edges represent paths from SS18-SSX to p21. **(d)** Model of regulation and intervention in the core oncogenic program. SS18-SSX activates the core oncogenic program in an HDAC-dependent manner and promotes cell cycle through direct activation of *CDK6* and *CCND2* (CycD) transcription. The core program suppresses p21 and inhibits immunogenic features. HDAC and CDK6 inhibitors target SyS dependencies.

To test these predictions, we treated primary SyS cell cultures with TNF and IFN*γ*, separately and in combination, and profiled 1,050 cells by scRNA-Seq. As predicted, combined TNF and IFN*γ* treatment repressed the core oncogenic program (P = 6.66*10^-18^, mixed-effects, **Fig. 6B**) in a synergistic manner (P = 9.49*10^-4^, interaction term, mixed-effects). Moreover, the treatment repressed the predicted TNF/IFN*γ*-dependent component of the program (1.6*10^-38^, mixed-effects), but not the component predicted to be TNF/IFN*γ*-independent (P > 0.05, **Fig. 6B**). The combined treatment also repressed the SS18-SSX program (P < 3.12*10^-16^, both direct and indirect components, including *TLE1*; P = 1.23*10^-4^ for the interaction term, **Fig. 6B, Supp. Table 6B**), and induced multiple genes from the epithelial program (P = 1.95*10^-9^, hypergeometric test, **Supp. Table 6B**). Short-term (4-6 hours) treatment with TNF alone substantially repressed homeobox genes (*e.g., MEOX2*, **Supp. Table 6C**), which are directly bound by SS18-SSX (18,19) (P < 1*10^-17^, hypergeometric test). It also repressed the core oncogenic program, but only temporarily (P = 8.73*10^-18^, mixed-effects; **Supp. Fig. 5D**), suggesting that IFN*γ* is required to sustain the effect. Interestingly, TNF also induced TNF expression in the SyS cells (P < 5.57*10^-8^, mixed-effects), suggesting that autocrine signaling might induce the effect. Taken together, these findings demonstrate that macrophages and T cells can suppress the SS18-SSX program by secreting TNF and IFN*γ*.

### HDAC and CDK4/6 inhibitors synergistically repress the immune resistant features of synovial sarcoma cells

Lastly, we turned to examine whether pharmacological agents could potentially repress the core oncogenic program and induce more immunogenic cell states in SyS cells. Computational modeling of the core oncogenic regulatory network (**Methods**) highlighted the SSX-SS18-HDAC1 complex (20) as the program’s master regulator (**Fig. 6C**), and the tumor suppressor *CDKN1A* (p21) as its most repressed target. The latter indicates that the core oncogenic program is regulating, rather than regulated by, cell cycle genes through the p21-CDK2/4/6 axis, potentially reinforcing the direct induction of cyclin D and *CDK6* by SS18-SSX (**Fig. 6C,D**). According to this model (**Fig. 6D**), modulators of cell cycle (*e.g.*, CDK4/6 inhibitors) and SS18-SSX (*e.g.*, HDAC inhibitors) could synergistically target the immune resistance features of SyS cells, especially in the presence of tumor microenvironment cytokines as TNF.

To test these predictions, we treated SyS lines and primary mesenchymal stem cells (MSCs) with low doses of HDAC and CDK4/6 inhibitors, in order to avoid global toxicity-related effects, and examined their impact on the transcriptional state of the cells. As predicted, the HDAC inhibitor panobinostat markedly repressed the core oncogenic program (P = 3.34*10^-14^, mixed-effects; **Fig. 7A**) and selectively induced *CDKN1A* in SyS cells (P = 2.13*10^-8^) (**Supp. Fig. 6A**). Panobinostat also repressed the SS18-SSX program (P = 5.32*10^-72^; **Fig. 7B**), decreased the expression of cell cycle genes (P < 1.78*10^-20^), and induced an immunogenic phenotype (32) with enhanced antigen presentation and IFN*γ* responses (P < 9.53*10^-31^; **Fig. 7C,D, Supp. Fig. 6B,C**). The CDK4/6 inhibitor abemaciclib repressed cell cycle gene expression (P = 3.63*10^-8^), without impacting the core oncogenic program (P > 0.1; **Fig. 7A**), supporting the notion that cell cycle regulation is down-stream of the core oncogenic program. Lastly, a low dose combination of panobinostat, abemaciclib and TNF synergistically repressed the core oncogenic program (P = 1.72*10^-37^, **Fig. 7A, Supp. Fig. 6A**) and multiple immune resistant features, while inducing antigen presentation, IFN responses, and induced-self antigens as MICA/B (P = 3.12*10^-76^; **Fig. 7C,D, Supp. Fig. 6B,C**). It also repressed *MIF* (Macrophage Migration Inhibitory Factor), a member of the core oncogenic and SS18-SSX programs, which has been previously shown to hamper T cell recruitment into the tumor (40). The effect of the drug combination on these programs and genes in viable SyS cells significantly exceeded the expected additive effect (P < 0.01, mixed-effects interaction term, **Methods**), and could potentially help both T cells (MHC-1) and NK cells (MICA/B) bind to and eliminate SyS cell. Consistent with the transcriptional changes, the drug combination displayed a significantly higher detrimental effect on the SyS cells compared to primary MSCs (P = 5.7*10^-13^; **Fig. 7E,F**).

**Figure 7.**
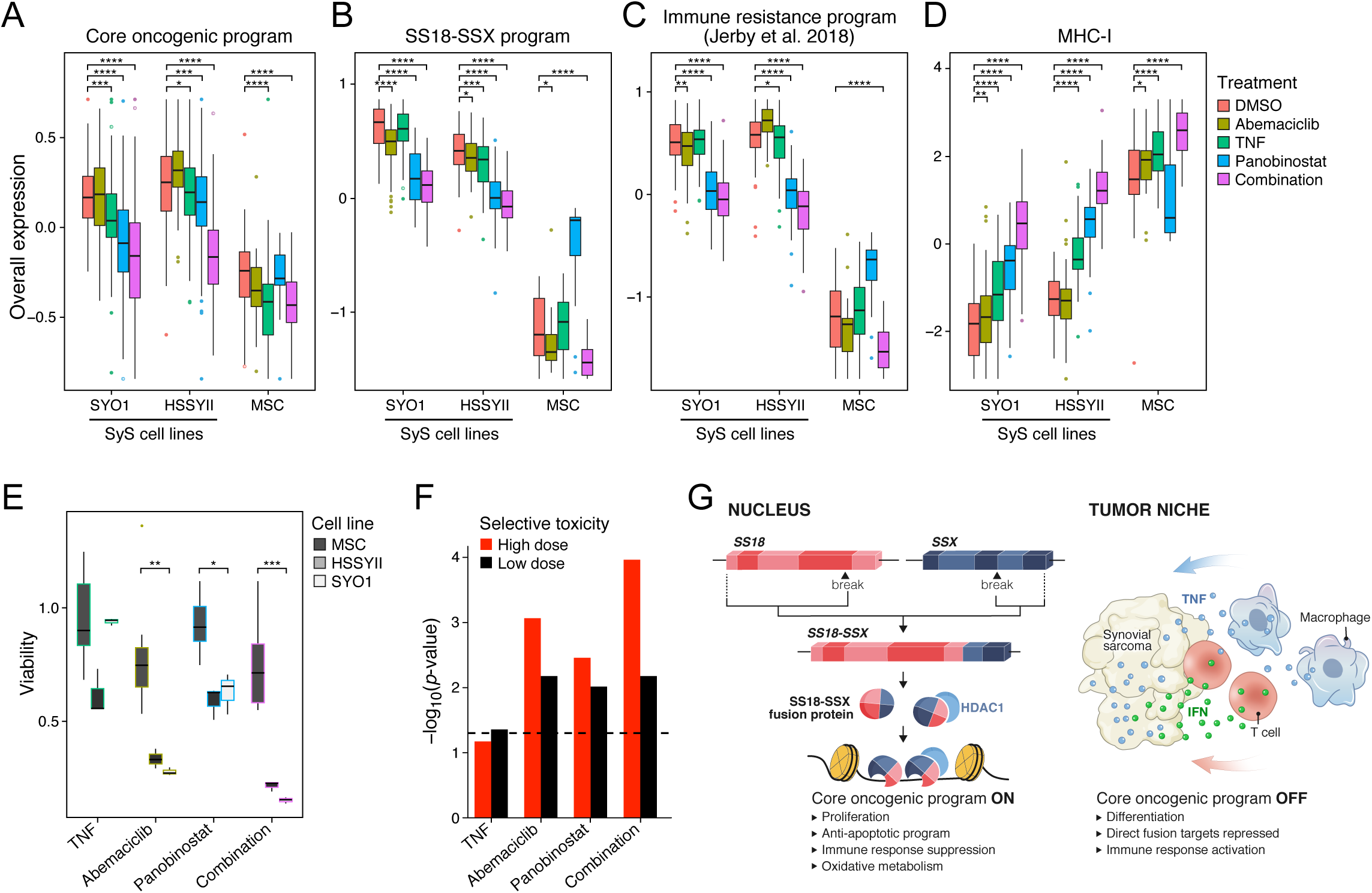
The core oncogenic program can be selectively blocked in SyS cells by combined HDAC and CDK4/6 inhibitors. **(a-d)** TNF, HDAC and CDK6 inhibitors suppress the core oncogenic program. Overall Expression of the core oncogenic program **(a)**, SS18-SSX program **(b)**, an immune resistance program identified in melanoma (32) **(c)**, and MHC-1 genes **(d)** in SyS cells and MSCs (x axis). (A-D) *P<0.1,**P<0.01, ***P<1*10^-3^, ****P<1*10^-4^, t-test. **(e,f)** Selective toxicity for SyS cell lines. (**e**) Viability (*y* axis) of SyS cell lines and MSCs (*x* axis) under different drugs (*x* axis, *P<5*10^-2^, **P<5*10^-3^, ***P<5*10^-4^, ANOVA test). **(f)** Selective toxicity to SyS lines *vs*. MSC (y axis, -log_10_(P-value), ANOVA) in each treatment (x axis). In (A-E) middle line: median; box edges: 25_th_ and 75_th_ percentiles, whiskers: most extreme points that do not exceed ±IQR*1.5; further outliers are marked individually. **(g)** Model of intrinsic and microenvironment determinants of SyS cell states. Left: The SS18-SSX oncoprotein sustains de-differentiation, proliferation and the core oncogenic program. Right: immune cells in the tumor microenvironment can repress the core oncogenic and SS18-SSX programs through TNF and IFN*γ* secretion. Combined inhibition of HDAC and CDK4/6 mimics these effects selectively in SyS cells.

## DISCUSSION

Here, we mapped malignant and immune cell states and interactions in human SyS tumors, through integrative analyses of clinical and functional data. Out data reveals active antitumor immunity in this relatively cold tumor, alongside malignant cellular plasticity and immune excluding features, centered around a core oncogenic program. This program is regulated by the tumor’s primary genetic driver and may hamper proper immune recruitment and infiltration. Nonetheless, immune cells can impact the malignant cells through TNF and IFN*γ* secretion, counteracting the transcriptional alterations induced by the oncoprotein. Targeting the oncogenic program and its downstream effects with HDAC and CDK4/6 inhibitors induced cell autonomous immune responses, repressed immune resistant features, and was selectively detrimental to SyS cells.

The metabolic features of the core oncogenic program may also impact the tumor microenvironment. Supporting this notion, recent studies have shown that malignant cells use oxidative phosphorylation to create a hypoxic niche and promote T cell dysfunction (41). These metabolic features might reflect the conserved role of the SWI/SNF complex in regulating carbon metabolism and sucrose non-fermenting phenotypes in the yeast *Saccharomyces cerevisiae* (42). These connections might generalize to other cancer types, as mutations in the BAF complex have been recently shown to induce a targetable dependency on oxidative phosphorylation in lung cancer (43).

Despite the extremely cold phenotypes displayed by SyS (**Fig. 2D**), expanded effector T cells are present in SyS tumors (**Fig. 2B,C**), potentially responding to the CTAs expressed specifically in the malignant cells, including NY-ESO-1 and *PRAME* (**Supp. Fig. 2A**). Consistently, vaccines triggering dendritic-cells to prime NY-ESO-1 specific T cells can lead to durable responses in SyS patients (7), further supporting the notion that SyS immune evasion operates primarily through impaired T cell or dendritic cell recruitment (44). The latter may also be mediated through Wnt/β-catenin signaling pathway, which has been previously shown to interfere with CD8 T cell recruitment to tumors by dendritic cells (44), and is indeed active in all the malignant SyS cells and directly induced by SS18-SSX (**Supp. Fig. 5B, Supplementary Tables 2, 6**). The core oncogenic program itself includes several CTAs, linking between malignant immune evasion and testicular immune privileges.

While the core oncogenic program shares some similar features with a T cell exclusion program we recently identified in melanoma (32), there are also substantial distinctions between the two programs, and >90% do not overlap between the two, likely reflecting the dramatic differences in driving events, cell of origin and tissue environment of the two tumors. This emphasizes the importance of understanding immune evasion for each tumor context. In particular, unlike the melanoma program, the core oncogenic program highlights a metabolic shift and is strongly connected to the genetic driver. In SyS tumors (but not in melanoma) we successfully decoupled, through computational inference, the intrinsic and extrinsic signals which modulate this transcriptional program, facilitating the reconstruction of multicellular circuits. This new approach revealed a bi-directional interaction between malignant and immune cells where CD8 T cells and macrophages can in turn repress the core oncogenic program through the secretion of TNF and IFN*γ*. Thus, beyond their direct cytotoxic activity, immune cells can alleviate some of the aggressive features of SyS cells through cytokine secretion, targeting also malignant cells with repressed antigen presentation or unrecognized epitopes.

Our findings also demonstrate that immune resistance, metabolic processes, cell cycle and de-differentiation are tightly co-regulated in SyS. Hence, certain targeted therapies may be able to sensitize the tumor to immune surveillance. Supporting this notion, we demonstrate that the combined inhibition of HDAC and CDK4/6, two known repressors of SS18-SSX (45,46) and cellular proliferation (47), respectively, trigger immunogenic cell states even at sub-cytotoxic doses. This combinatorial treatment is also selectively cytotoxic to SyS cells, consistent with previous reports where HDAC and CDK4/6 inhibitors were used separately to induce cell death in SyS (45,47). The basal antitumor immune response we report, and the ability of T cells and macrophages to repress the core oncogenic and SS18-SSX programs support the potential of exploiting HDAC and CDK4/6 inhibitors together with immunotherapy.

Taken together, our study comprehensively maps and interrogates cell states in SyS and its surrounding tumor microenvironment, along with their multicellular regulatory circuits and clinical implications. We demonstrated that the SS18-SSX oncoprotein and the tumor microenvironment coordinately shape cell states in SyS, resulting in the establishment of an immune privileged environment (**Fig. 7G**). The possibility to selectively target the underlying mechanisms to reverse immune evasion offers a new perspective for the clinical management of SyS, and potentially other malignancies driven by similar genetic events.

## ACKNOWLEDGMENTS

We thank Leslie Gaffney and Anna Hupalowska for help with artwork and Leslie Gaffney for help in figure preparation. We thank Kazuyuki Itoh, Norifumi Naka, and Satoshi Takenaka (Osaka University, Japan) for providing the Aska cell lines, and Akira Kawai (National Cancer Center Hospital, Japan) for providing the SYO1 cell line. L.J.A. is a CRI Irvington Fellow supported by the CRI, and a fellow of the Eric and Wendy Schmidt Postdoctoral program. Av.R. is an HHMI Investigator. Work was supported by the Klarman Cell Observatory, STARR cancer consortium, NCI grants 1U24CA180922, R33-CA202820, the Koch Institute NCI Support (core) grant P30-CA14051, Ludwig Centers at Harvard and MIT, AMRF and the Broad Institute (Av.R.). Work was also supported by grants from the Howard Goodman Fellowship at MGH (M.L.S.), the Merkin Institute Fellowship at the Broad Institute of MIT and Harvard (M.L.S.), and the Swiss National Science Foundation Sinergia grant (M.L.S. and I.S.). Imaging CyCIF work was supported by a grant (CA225088) from the Center for Cancer Systems Pharmacology at Harvard Medical School (P.K.S.). N.R. is supported by the Swiss National Science Foundation Professorship grant (PP00P3-157468/1 and PP00P3_183724), the Swiss Cancer League grant KFS-3973-08-2016, the Fond’Action Contre le Cancer grant and the FORCE grant. Av.R. is a scientific advisory board member for ThermoFisher Scientific, Syros Pharmaceuticals and Driver Group. L.J.A, N.R., M.L.S. and Av.R. are co-inventors on US patent application filed by the Broad Institute relating to synovial sarcoma. D.R.Z. and J.M.B. are employees of Nanostring which developed GeoMx. Processed scRNA-Seq data will be available through the Gene Expression Omnibus (GEO).

## AUTHOR CONTRIBUTION

L.J.A., C.N., N.R., M.L.S. and Av.R., conceived the project, designed the study, and interpreted results. Av.R., O.R.R., N.R., and M.L.S. obtained funding for the study. L.J.A. performed computational analyses. C.N., M.E.S., H.R.W., A.R.R., G.B., A.V., collected synovial sarcoma samples and generated single-cells RNA-sequencing data. B.I. D.R.Z. and J.M.B. performed tissue spatial analyses. B.H. provided support for single-cell genetic analyses. C.C.L. and R.M. provided flow cytometry expertise. M.J.M. and C.K. provided data and support for chromatin analysis. G.P.N., I.C., G.C., E.C., S.C., P.K.S., A.B.H., J.T.M., consented patients for the study and provided clinical data. I.L., L.C., L.C.B, J.M.G., L.N., S.M., J.C.M., C.G., O.C., J.B., M.S.C., D.L., N.W., I.S., M.N.R. and O.R.R. provided experimental and analytical support. M.L.S., N.R. and Av.R. jointly supervised this work. L.J.A, N.R., M.L.S. and Av.R. wrote the manuscript with feedback from all authors.

## METHODS

### Human tumor specimen collection and dissociation

Patients at Massachusetts General Hospital and University Hospital of Lausanne were consented preoperatively in all cases according to their respective Institutional Review Boards (protocol numbers: CER-VD 260/15, DF/HCC 13-416). Fresh tumors were collected directly from the operating room at the time of surgery and presence of malignancy was confirmed by frozen section. Tumor tissues were mechanically and enzymatically dissociated using a human tumor dissociation kit (Miltenyi Biotec, Cat. No. 130-095-929), following the manufacturers recommendations. Clinical annotations are provided in **Supp. Table 1**.

### Fluorescence-activated cell sorting (FACS)

Tumor cells were kept in Phosphate Buffered Saline with 1% bovine serum albumin (PBS/BSA) while staining. Cells were stained using calcein AM (Life Technologies) and TO-PRO-3 iodide (Life Technologies) to identify viable cells. For all tumors, we used CD45-VioBlue (human antibody, clone REA747, Miltenyi Biotec) to identify immune cells and in few cases, we also used CD3-PE to specifically identify lymphocytes (human antibody, clone BW264/56, Miltenyi Biotec). For all the samples, we used unstained cells as control. Standard, strict forward scatter height versus area criteria were used to discriminate doublets and gate only single cells. Viable single cells were identified as calcein AM positive and TO-PRO-3 negative. Sorting was performed with the FACS Aria Fusion Special Order System (Becton Dickinson) using 488nm (calcein AM, 530/30 filter), 640nm (TO-PRO-3, 670/14 filter), 405nm (CD45-VioBlue, 450/50 filter) and 561nm (PE, 586/15 filter) lasers. We sorted individual, viable, immune and non-immune single cells into 96-well plates containing TCL buffer (Qiagen) with 1% beta-mercaptoethanol. Plates were snap frozen on dry ice right after sorting and stored at −80°C prior to whole transcriptome amplification, library preparation and sequencing.

### Library construction and sequencing

For plate-based scRNA-seq, Whole transcriptome amplification was performed using the SMART-seq2 protocol (24), with some modifications as previously described (30,49,50). The Nextera XT Library Prep kit (Illumina) was used for library preparation, with custom barcode adapters (sequences available upon request). Libraries from 384 to 768 cells with unique barcodes were combined and sequenced using a NextSeq 500 sequencer (Illumina).

In addition to SMART-seq2, cells from three samples (SS12pT, SS13 and SS14) were also sequenced using droplet-based scRNA-Seq with the 10x genomics platform. The samples were partitioned for SMART-seq2 and 10x genomics after dissociation. For each tumor, approximately two thirds of the sample was used for SMART-seq2 and one third for droplet based scRNA-seq (10x genomics). We sorted viable cells using MACS (Dead Cell Removal Kit, Miltenyi Biotec) and ran up to 2 channels per sample with a targeted number of cell recovery of 2,000 cells per channel. The samples were processed using the 10x Genomics Chromium 3’ Gene Expression Solution (version 2) based on manufacturer instructions and sequenced using a NextSeq 500 sequencer (Illumina).

### Whole exome sequencing (WES)

DNA and RNA were extracted from fresh frozen tissue or Formalin-Fixed Paraffin-Embedded (FFPE) blocks for each patient (obtained according to their respective Institutional Review Board-approved protocols) using the AllPrep DNA/RNA extraction kit (Qiagen). We used tumor tissue and matched normal muscle tissue from the same patient as reference. Library construction was performed as previously described (50), with the following modifications: initial genomic DNA input into shearing was reduced from 3µg to 20-250ng in 50µL of solution. For adapter ligation, Illumina paired end adapters were replaced with palindromic forked adapters, purchased from Integrated DNA Technologies, with unique dual-indexed molecular barcode sequences to facilitate downstream pooling. Kapa HyperPrep reagents in 96-reaction kit format were used for end repair/A-tailing, adapter ligation, and library enrichment PCR. In addition, during the post-enrichment SPRI cleanup, elution volume was reduced to 30µL to maximize library concentration, and a vortexing step was added to maximize the amount of template eluted. After library construction, libraries were pooled into groups of up to 96 samples. Hybridization and capture were performed using the relevant components of Illumina’s Nextera Exome Kit and following the manufacturer’s suggested protocol, with the following exceptions: first, all libraries within a library construction plate were pooled prior to hybridization. Second, the Midi plate from Illumina’s Nextera Exome Kit was replaced with a skirted PCR plate to facilitate automation. All hybridization and capture steps were automated on the Agilent Bravo liquid handling system. After post-capture enrichment, library pools were quantified using qPCR (automated assay on the Agilent Bravo), using a kit purchased from KAPA Biosystems with probes specific to the ends of the adapters. Based on qPCR quantification, libraries were normalized to 2nM. Cluster amplification of DNA libraries was performed according to the manufacturer’s protocol (Illumina), using exclusion amplification chemistry and flowcells. Flowcells were sequenced using Sequencing-by-Synthesis chemistry. The flowcells are then analyzed using RTA v.2.7.3 or later. Each pool of whole exome libraries was sequenced on paired 76 cycle runs with two 8 cycle index reads across the number of lanes needed to meet coverage for all libraries in the pool.

### *In situ* immunofluorescence imaging

Formalin-fixed, paraffin-embedded (FFPE) tissue slides, 5 µm in thickness, were generated at the at the Massachusetts General Hospital from tissue blocks collected from patients under IRB-approved protocols (DF/HCC 13-416). Multiplexed, tissue cyclic immunofluorescence (t-CyCIF) was performed as described recently (51). For direct immunofluorescence, we used the following antibodies (manufacturer, clone, dilution): c-Jun-Alexa-488 (Abcam, Clone E254, 1:200), CD45-PE (R&D, Clone 2D1, 1:150), p21-Alexa-647 (CST, Clone 12D1, 1:200), Hes1-Alexa-488 (Abcam, Clone EPR4226, 1:500), FoxP3-Alexa-570 (eBioscience, Clone 236A/E7, 1:150), NF-κB (Abcam, Clone E379, 1:200), E-Cadherin-Alexa-488 (CST, Clone 24E10, 1:400), pRB-Alexa-555 (CST, Clone D20B12, 1:300), COXIV-Alexa-647 (CST, Clone 3E11, 1:300), β-catenin-Aleaxa-488 (CST, Clone L54E2, 1:400), HSP90-PE (Abcam, polyclonal, lot# GR3201402-2, 1:500) and vimentin-Alexa-647 (CST, Clone D21H3, 1:200). Stained slides from each round of t-CyCIF were imaged with a CyteFinder slide scanning fluorescence microscope (RareCyte Inc. Seattle WA) using either a 10X (NA=0.3) or 40X long-working distance objective (NA = 0.6). Imager5 software (RareCyte Inc.) was used to sequentially scan the region of interest in 4 fluorescence channels. Image processing, background subtraction, image registration, single-cell segmentation and quantification were performed as previously described (51).

### RNA *in situ* hybridization

Paraffin-embedded tissue sections from human tumors from Massachusetts General Hospital and and University Hospital of Lausanne were obtained according to their respective Institutional Review Board-approved protocols. Sections were mounted on glass slides and stored at −80°C. Slides were stained using the RNAscope 2.5 HD Duplex Detection Kit (Advanced Cell Technologies, Cat. No. 322430), as previously described (30,49,52): slides were baked for 1 hour at 60°C, deparaffinized and dehydrated with xylene and ethanol. The tissue was pretreated with RNAscope Hydrogen Peroxide (Cat. No. 322335) for 10 minutes at room temperature and RNAscope Target Retrieval Reagent (Cat. No. 322000) for 15 minutes at 98°C. RNAscope Protease Plus (Cat. No. 322331) was then applied to the tissue for 30 minutes at 40°C. Hybridization probes were prepared by diluting the C2 probe (red) 1:50 into the C1 probe (green). Advanced Cell Technologies RNAscope Target Probes used included Hs-EGR1 (Cat. No. 457671-C2) and Hs-IGF2 (Cat. No. 594361). Probes were added to the tissue and hybridized for 2 hours at 40°C. A series of 10 amplification steps was performed using instructions and reagents provided in the RNAscope 2.5 HD Duplex Detection Kit. Tissue was counterstained with Gill’s hematoxylin for 25 seconds at room temperature followed by mounting with VectaMount mounting media (Vector Laboratories).

### RNA profiling in situ hybridization (ISH)

DNA oligo probes were designed to bind mRNA targets. From 5’ to 3’, they each comprised of a 35-50 nt target complementary sequence, a UV photocleavable linker, and a 66 nt indexing oligo sequence containing a unique molecular identifier (UMI), RNA ID sequence, and primer binding sites. Up to 10 RNA detection probes were designed per target mRNA. RNA detection probes were provided by Nanostring Technologies.

To perform the ISH, 5 µm FFPE tissue sections from two patients were mounted on positively charged histology slides. Sections were baked at 65°C for 45 minutes in a Hyb EZ II hybridization oven (Advanced cell Diagnostics, Inc). Slides were deparaffinized using Citrsolv (Decon Labs, Inc., 1601), rehydrated in an ethanol gradient, and washed in 1x phosphate-buffered saline pH 7.4 (PBS: Invitrogen, AM9625). Slides were incubated for 15 minutes in 1X Tris-EDTA pH 9.0 buffer (Sigma Aldrich, SRE0063) at 100°C with low pressure in a TintoRetriever Pressure cooker (bioSB, 7008). Slides were washed, then incubated in 1 µg/mL proteinase K (Thermo Fisher Scientific, AM2546) in PBS for 15 minutes at 37°C and washed again in PBS. Tissues were then fixed in 10% neutral-buffered formalin (Thermo Fisher Scientific, 15740) for 5 minutes, incubated in NBF stop buffer (0.1M Tris Base, 0.1M Glycine, Sigma) for 5 minutes twice, then washed for 5 minutes in PBS. Tissues were then incubated overnight at 37°C with GeoMx™ RNA detection probes in Buffer R (Nanostring Technologies) using a Hyb EZ II hybridization oven (Advanced cell Diagnostics, Inc). During incubation, slides were covered with HybriSlip Hybridization Covers (Grace BioLabs, 714022). Following incubation, HybriSlip covers were gently removed and 25-minute stringent washes were performed twice in 50% formamide and 2X SSC at 37°C. Tissues were washed for 5 minutes in 2X SSC then blocked in Buffer W (Nanostring Technologies) for 30 minutes at room temperature in a humidity chamber. 500nM Syto13 and antibodies targeting PanCK and CD45 (Nanostring technologies) in Buffer W were applied to each section for 1 hour at room temperature. Slides were washed twice in fresh 2X SSC then loaded on the GeoMx™ Digital Spatial Profiler (DSP) (53). In brief, entire slides were imaged at 20x magnification and 12 circular regions of interest (ROI) with 200-300 μm diameter were selected per sample. The DSP then exposed ROIs to 385 nm light (UV) releasing the indexing oligos and collecting them with a microcapillary. Indexing oligos were then deposited in a 96-well plate for subsequent processing. The indexing oligos were dried down overnight and resuspended in 10 μL of DEPC-treated water.

Sequencing libraries were generated by PCR from the photo-released indexing oligos and ROI-specific Illumina adapter sequences and unique i5 and i7 sample indices were added. Each PCR reaction used 4 μL of indexing oligos, 1 μL of indexing PCR primers, 2 μL of Nanostring 5X PCR Master Mix, and 3 μL PCR-grade water. Thermocycling conditions were 37°C for 30 min, 50°C for 10 min, 95°C for 3 min; 18 cycles of 95°C for 15sec, 65°C for 1min, 68°C for 30 sec; and 68°C 5 min. PCR reactions were pooled and purified twice using AMPure XP beads (Beckman Coulter, A63881) according to manufacturer’s protocol. Pooled libraries were sequenced at 2×75 base pairs and with the single-index workflow on an Illumina NextSeq to generate 458M raw reads.

### Primary cell cultures and cell lines

Human primary Synovial Sarcoma (SyS) spherogenic cultures (SScul1, SScul2 and SScul3) were derived from patients undergoing surgery at Massachusetts General Hospital and University Hospital of Lausanne, according to their respective Institutional Review Board-approved protocols. Directly after dissociation (as above), the dissociated bulk tumor cells were put in culture and grown as spheres using ultra-low attachment cell culture flasks in IMDM 80% (Gibco, Cat. No. 1244053), Knock-Out Serum Replacement 20% (Gibco, Cat. No. 10828028), Recombinant Human EGF Protein 10 ng/mL (R&D systems, Cat. No. 236-EG-200), Recombinant Human FGF basic, 145 aa (TC Grade) Protein 10ng/mL (R&D systems, Cat. No. 4114-TC-01M) and 1% Penicillin-Streptomycin (Gibco, Cat. No. 15140122). Cells were expanded by mechanical and enzymatical dissociation every week using TrypLE Express Enzyme (ThermoFisher, Cat. No. 12605010).

The SyS cell lines used for the SS18-SSX KD experiments and the functional drug assays include: Aska (a generous gift from Kazuyuki Itoh, Norifumi Naka, and Satoshi Takenaka, Osaka University, Japan), SYO1 (a generous gift from Akira Kawai, National Cancer Center Hospital, Japan), and HS-SY-II (purchased from RIKEN Bio Resource Center, 3-1-1 Koyadai, Tsukuba, Ibaraki 305-0074, Japan). All three cell lines were cultured using standard protocols in DMEM medium (Gibco) supplemented with 10-20% fetal bovine serum, 1% Glutamax (Gibco), 1% Sodium Pyruvate (Gibco) and 1% Penicillin-Streptomycin (Gibco) and grown in a humidified incubator at 37°C with 5% CO_2_.

Human primary pediatric Mesenchymal Stem Cells (MSCs) were isolated from healthy donors undergoing corrective surgery in agreement with the Institutional Review Board-approved protocol of the University Hospital of Lausanne (Protocol number 2017-0100). Samples were deidentified prior to culture and analysis. Cells were expanded in 90% IMDM (Gibco, Cat. No. 1244053) containing 10% Fetal Bovine Serum (Gibco), 1% Penicillin-Streptomycin (Gibco) and 10ng/mL Platelet-Derived Growth Factor BB (PDGF-BB, PeproTech) as previously described (54).

### SS18-SSX knockdown in Aska and SYO1 cell lines

The SyS cell lines Aska and SYO1 were cultured using standard protocols in DMEM medium (Gibco) supplemented with 10-20% fetal bovine serum, 1% Glutamax (Gibco), 1% Sodium Pyruvate (Gibco) and 1% Penicillin-Streptomycin (Gibco) and grown in a humidified incubator at 37°C with 5% CO_2_. Cells expressing a pLKO.1 vector with a scrambled shRNA hairpin control (5’-CCTAAGGTTAAGTCGCCCTCGCTCGAGCGAGGGCGACTTAAC CTTAGG-3’) or a shSSX hairpin targeting SSX of the SS18-SSX fusion (5’-CAGTCACTGACAGTTAATAAA-3’) were prepared by lentiviral infection. Briefly, lentivirus was prepared by transfection of HEK293T cells with gene delivery vector and the packaging vectors pspax2 and pMD2.G, filtration of media followed by ultracentrifugation, and then resuspension of viral pellet in PBS. Aska and SYO1 cells were infected with lentivirus for 48 hours and then underwent 5 days of selection with puromycin (2 μg/mL) prior to collection for scRNA-seq.

### *In vitro* IFN/TNF experiment

Cells were dissociated 12 hours before adding the drugs at the concentrations indicated directly to the growing media and cells were collected at different time point (ranging from 4 hours to 4 days) for SMART-seq2. Viability was determined by CellTiter-Glo Luminescent Cell Viability Assay (Promega) after 5 to 7 days of treatment. TNF-alpha (Miltenyi Biotec, Human TNF-α, Cat. No. 130-094-014) IFN-gamma (R&D systems, Recombinant Human IFN-gamma Protein, Cat. No. 285-IF-100) were suspended in deionized sterile-filtered water.

### *In vitro* drug assay and cell proliferation measurements

For the functional drug assay, 200,000 SYO-1 cells and HSSYII cells, and 100,000 MSCs were seeded in 60 × 15 mm plates (Falcon). Cells were stimulated for five days with the following compounds: 100 or 200 nM Abemaciclib (Selleckchem, U.S.A.), 15 or 30 ng/ml TNF (Miltenyi Biotech, Germany) or a combination of the two. Compounds were refreshed at days three and four, and the solvent (DMSO) was used as control. At day 4, 12.5 or 25 nM Panobinostat (Selleckchem, U.S.A.) was added to the cultures, and the cells were harvested 24 hours later for proliferation scoring. To assessment cellular proliferation, cells were detached with trypsin, washed in PBS, and re-suspended in 1 ml of complete medium. After diluting 1:2 with Trypan blue (Invitrogen) viable cells were counted using the Automated Cell Counter Countess II FL (Thermo Fisher Scientific). Each experimental condition was measured in triplicate.

### scRNA-seq pre-processing and gene expression quantification

BAM files were converted to merged, demultiplexed FASTQ files. The paired-end reads obtained with SMART-Seq2 were mapped to the UCSC hg19 human transcriptome using Bowtie (55), and transcript-per-million (TPM) values were calculated with RSEM v1.2.8 in paired-end mode (56). The paired-end reads obtained with droplet scRNA-Seq (10x Genomics) were mapped to the UCSC hg19 human transcriptome using STAR (57), and gene counts/TPM values were obtained using CellRanger (cellranger-2.1.0, 10x Genomics).

For bulk RNA-Seq, expression levels were quantified as E=log_2_(TPM+1). For scRNA-seq data, expression levels were quantified as E=log_2_(TPM_i,j_/10+1). TPM values were divided by 10 because the complexity of our single-cell libraries is estimated to be within the order of 100,000 transcripts (52). The 10^-1^ factoring prevents counting each transcript ∼10 times and overestimating differences between positive and zero TPM values. The average expression of a gene *i* across a population *P* of *N* cells, was defined as

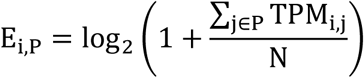

For each cell, we quantified the number of genes with at least one mapped read, and the average expression level of a curated list of housekeeping genes (58). We excluded all cells with either fewer than 1,700 detected genes or an average housekeeping expression (*E*, as defined above) below 3 (**Supp. Table 1**). For the remaining cells, we calculated the average expression of each gene (*E*_*p*_), and excluded genes with an average expression below 4, which defined a different set of genes in different analyses depending on the subset of cells included. In cases where we analyzed different cell subsets together, we removed genes only if they had an average E_p_ below 4 in each of the different cell subsets included in the analysis. Different cell types and malignant cells from different tumors were considered as different cell subsets in this regard.

### WES data pre-processing

A BAM file was produced with the Picard pipeline (http://picard.sourceforge.net/), which aligns the tumor and normal sequences to the hg19 human genome build using Illumina sequencing reads. The BAM was uploaded into the Firehose pipeline (http://www.broadinstitute.org/cancer/cga/Firehose). Quality control modules within Firehose were applied to all sequencing data for comparison of the origin for tumor and normal genotypes and to assess fingerprinting concordance. Cross-contamination of samples was estimated using ContEst (59).

### Somatic alteration assessment

MuTect (60) was applied to identify somatic single-nucleotide variants. Indelocator (http://www.broadinstitute.org/cancer/cga/indelocator), Strelka (61), and MuTect2 (https://software.broadinstitute.org/gatk/documentation/tooldocs/current/org_broadinstitute_gatk_tools_walkers_cancer_m2_MuTect2) were applied to identify small insertions or deletions. A voting scheme was used with inferred indels requiring a call by at least 2 out of 3 algorithms.

Artifacts introduced by DNA oxidation during sequencing were computationally removed using a filter-based method (62). In the analysis of primary tumors that are formalin-fixed, paraffin-embedded samples (FFPE) we further applied a filter to remove FFPE-related artifacts (63). Reads around mutated sites were realigned with Novoalign (www.novocraft.com/products/novoalign/) to filter out false positive that are due to regions of low reliability in the reads alignment. At the last step, we filtered mutations that are present in a comprehensive WES panel of 8,334 normal samples (using the Agilent technology for WES capture) aiming to filter either germline sites or recurrent artifactual sites. We further used a smaller WES panel of 355 normal samples that are based on Illumina technology for WES capture, and another panel of 140 normal samples sequenced without our cohort (64) to further capture possible batch-specific artifacts. Annotation of identified variants was done using Oncotator (65) (http://www.broadinstitute.org/cancer/cga/oncotator).

### Copy number and copy ratio analysis

To infer somatic copy number from WES, we used ReCapSeg (http://gatkforums.broadinstitute.org/categories/recapseg-documentation), calculating proportional coverage for each target region (*i.e.*, reads in the target/total reads) followed by segment normalization using the median coverage in a panel of normal samples. The resulting copy ratios were segmented using the circular binary segmentation algorithm (66). To infer allele-specific copy ratios, we mapped all germline heterozygous sites in the germline normal sample using GATK Haplotype Caller (67) and then evaluated the read counts at the germline heterozygous sites in order to assess the copy profile of each homologous chromosome. The allele-specific copy profiles were segmented to produce allele specific copy ratios.

### Gene sets Overall Expression

We used the following scheme to compute the Overall Expression (OE) of a gene set (signature). The OE metric (32) filters technical variation and highlights biologically meaningful patterns. The procedure is based on the notion that the measured expression of a specific gene is correlated with its true expression (signal), but also contains a technical (noise) component. The latter may be due to various stochastic processes in the capture and amplification of the gene’s transcripts, sample quality, as well as variation in sequencing depth. The OE of a gene signature is computed in a way that accounts for the variation in the signal-to-noise ratio across genes and cells.

Given a gene signature and a gene expression matrix *E* (as defined above), we first binned the genes into 50 expression bins according to their average expression across the cells or samples. The average expression of a gene across a set of cells within a sample is *E*_*i,p*_ (see: **scRNA-seq pre-processing and gene expression quantification**) and the average expression of a gene across a set of *N* tumor samples was defined as 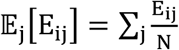. Given a gene signature *S* that consists of *K* genes, with *k*_*b*_ genes in bin *b*, we sample random *S-compatible* signatures for normalization. A random signature is *S-compatible* with a signature *S* if it consists of overall *K* genes, such that in each bin *b* it has exactly *k*_*b*_ genes. The OE of signature *S* in cell or sample *j* is then defined as:

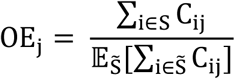

Where 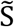 is a random S-compatible signature, and C_ij_ is the centered expression of gene *i* in cell or sample *j*, defined as C_ij_ = E_ij_ − 𝔼[E_ij_]. Because the computation is based on the centered gene expression matrix *C*, genes that generally have a higher expression compared to other genes will not skew or dominate the signal. We found that 100 random S-compatible signatures are sufficient to yield a robust estimate of the expected value 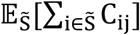. The distribution of the OE values was normal or a mixture of normal distributions, facilitating subsequent analyses.

We use the term *transcriptional program* (*e.g.*, the core oncogenic program) to denote cell states defined by a pair of signatures, such that one (S-up) is overexpressed and the other (S-down) is underexpressed. The OE of a program is then the OE of S-up minus the OE of S-down.

In cases where the OE of a given signature/program has a bimodal distribution across the cell population, it can be used to naturally separate the cells into two subsets. To this end, we applied the Expectation Maximization (EM) algorithm for mixtures of normal distributions to define the two underlying normal distributions. We then assigned cells to two subsets, depending on the distribution (high or low) they were assigned to.

### Cell type assignments

Cell type assignments were performed based on genetic and transcriptional features, according to the following four analyses:

#### (1) Fusion detection

Fusion detection was performed with STAR-Fusion (26), to detect any transcript that indicates the fusion of two genes.

#### (2) Copy Number Alterations (CNA) inference

To infer CNAs from the scRNA-seq data we used the approach described in (58), as implemented in the R code provided in https://github.com/broadinstitute/inferCNA with the default parameters. To identify malignant cells based on CNA patterns, we defined the overall CNA level of a given cell as the sum of the absolute CNA estimates across all genomic windows. Within each tumor, we identified CD45^-^ cells with the highest overall CNA level (top 10%), and considered their average CNA profile as the CNA profile of the pertaining tumor. For each cell we then computed a CNA-R-score defined as the Spearman correlation coefficient obtained when comparing its CNA profile to the inferred CNA profile of its tumor. Cells with a high CNA-R-score (greater than the 25% of the CD45^-^ cell population) were considered as malignant according to the CNA criterion. As certain tumors/malignant cells have a stable genome, we did not use the CNA criterion to identify non-malignant cells. Large-scale CNAs were visualized (**Fig. 1G**) using a Bayesian approach, as described in https://github.com/broadinstitute/infercnv/wiki/infercnv-i6-HMM-type.

#### (3) Differential similarity to bulk tumors

We compared the scRNA-Seq profiles to those of bulk sarcoma tumors (23). RNA-Seq of bulk sarcoma tumors was downloaded from TCGA (http://xena.ucsc.edu). For each cell in our scRNA-Seq cohort we: (**i**) computed the Spearman correlation between its expression profile and the expression profiles of the bulk sarcoma tumors, and (**ii**) examined if the *r*_*s*_ coefficients obtained when comparing the cell to SyS tumors were higher than those obtained when comparing the cell to non-SyS sarcoma tumors, using a one-sided Wilcoxon ranksum test. Cells with a ranksum p-value < 0.05 were considered as potentially malignant, and as potentially non-malignant otherwise.

#### (4) Expression profile clustering

We clustered the cells by applying a shared nearest neighbor (SNN) modularity optimization algorithm (68), as implemented in the *Seurat* R package. First, Principle Component Analysis (PCA) was preformed and *k*-nearest neighbors (*k*NN) were calculated to construct the *k*-NN graph. The *k*-NN graph was used to identify clusters that optimize the modularity function. Next, we assigned clusters to cell types. Clusters where the majority of cells had the SS18-SSX1/2 fusion (by the method in (**1**)) were considered as malignant clusters. Non-malignant clusters were assigned to cell types by computing the OE of well-established cell type markers across the non-malignant cells (**Supp. Table 2**). The OE of each of these cell type signatures had a bimodal distribution across the cell population. Applying the Expectation Maximization (EM) algorithm for mixtures of normal distributions, we defined the two underlying normal distributions, and assigned cells to cell types. Each non-malignant cluster was enriched for cells of a particular cell type, and was assigned to that cell type.

We used these four converging criteria to assign the cells to nine cell subsets: malignant cells, epithelial cells, Cancer Associated Fibroblasts (CAFs), CD8 and CD4 T cells, B cells, NK cells, macrophages, and mastocytes. Specifically, a cell was labeled malignant if it was CD45^-^ and classified as malignant according to analyses (**3**) and (**4**) above. A cell was labeled non-malignant if it was classified as non-malignant according to analyses (**1-4**) above. Non-malignant cells were then further assigned to cell types based on their cluster assignment by (**4**). Cells with inconsistent assignments (157 in the SMART-Seq dataset and 558 in the droplet-based dataset) were removed from further analyses. Lastly, within malignant cells we identified epithelial cells by clustering each of the biphasic tumors into two clusters.

Cell type assignments were preformed separately for the SMART-Seq2 and droplet scRNA-Seq datasets cohort. Fusion detection was performed only with the full length SMART-Seq2 data.

### Cell type signatures

Cell type signatures were generated based on pairwise comparisons between identified cell subtypes: malignant cells, epithelial cells, CAFs, CD8 and CD4 T cells, B cells, NK cells, macrophages, and mastocytes. For each pair of cell subtypes we identified differentially expressed genes using the likelihood-ratio test (69), as implemented in the Seurat package (http://www.satijalab.org/seurat). Genes were considered as cell type specific if they were overexpressed in a particular cell subtype compared to all other cell subtypes (log-fold change > 0.25 and p-value < 0.05, following Bonferroni correction). We defined a general T cell signature for both CD4 and CD8 cells by identifying genes that were overexpressed in both CD4 and CD8 compared to all other (non T) cells. A more permissive version of this generic T cell signature includes genes which were overexpressed in CD4 *or* CD8 T cells compared to all other (non T) cells.

### Inferring tumor composition

Tumor composition was assessed based on the Overall Expression of the different cell type specific signatures we identified from the scRNA-seq data (**Supp. Table 2**). For example, the CD8 T cell signature was used to infer the level of CD8 T cells in the tumor, and likewise for other cell types. To estimate tumor purity we used the malignant SyS signature identified here (**Supp. Table 2**), which consists of genes that are exclusively expressed by malignant SyS cells compared to non-malignant cells in SyS tumors.

To evaluate the performance of this approach, we simulated 200 bulk RNA-Seq profiles. For each simulated bulk RNA-Seq profile we: (1) randomly chose one of the tumors in the cohort; (2) sampled 100 cells from different cell types profiled in this tumor – these cells include a mix of immune, stroma and malignant cells, at a randomly chosen composition; (3) summed the scRNA-Seq profiles of this randomly chosen population (*P*) of 100 cells, such that the bulk expression of gene *i* across this population was defined as

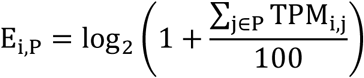

We also used cell type signatures we previously derived from melanoma scRNA-Seq data (32) to predict the tumor composition of the simulated SyS bulk RNA-Seq profiles, and vice versa. We then compared the predictions to the known cell type composition. The predicted composition was highly correlated with the known composition (*r* > 0.9, P < 1*10^-30^, Spearman correlation) for all cell types.

### Multilevel mixed-effects models

To examine the association between two cell features, denoted here as *x* and *y*, across different patients or experiments we used multilevel mixed-effects regression models (random intercepts models). The models include patient/experiment-specific intercepts to control for the dependency between the scRNA-seq profiles of cells that were obtained from the same patient/experiment. The models also control for data quality by providing the number of reads (log-transformed) that were detected in each cell as a covariate. To compute the association between features *x* and *y* we provided *x* as another covariate and used *y* as the dependent variable. The models were implemented using the lme4 and lmerTest R packages (https://CRAN.R-project.org/package=lme4, https://CRAN.R-project.org/package=lmerTest).

For example, to test if malignant cycling cells were more frequent in treatment naïve samples, we used a logistic mixed-effects model as described above. The dependent variable *y* was the cycling status of the malignant cells. The independent covariate *x* was a binary variable denoting if the sample was obtained before or after treatment. Only malignant cells were included in this model.

### T Cell Receptor (TCR) reconstruction and T cell expansion program

TCR reconstruction was performed using TraCeR (31), with the Python package in https://github.com/Teichlab/tracer. To characterize the transcriptional state of clonally expanded T cells, we first identified the clonality level of the T cells in our cohort. T cell that were obtained from tumors with a larger number of T cells with reconstructed TCRs were more likely to be defined as expanded. To control for this confounder we performed the following down-sampling procedure. First, we removed T cells without a reconstructed alpha or beta TCR chain, and samples with less than 20 T cells with a reconstructed TCR. Next, we computed the probability that a given cell will be a part of a clone when subsampling 20 T cells from each tumor. T cells with a high probability to be a part of a clone (above the median) were considered expanded, and non-expanded otherwise. To identify the genes differentially expressed in expanded CD8 T cells we used mixed-effects models with a binary covariate, denoting if the cell was classified as expanded or not.

### CD8 T cell analyses

The analysis of T cell exhaustion *vs*. T cell cytotoxicity was performed as previously described (58), with the exhaustion signature provided in (58). First, we computed the cytotoxicity and exhaustion scores of each CD8 T cell. Next, to control for the association between the expression of exhaustion and cytotoxicity markers, we estimated the relationship between the cytotoxicity and exhaustion scores using locally-weighted polynomial regression (LOWESS, black line in **Fig. 2B**). Based on these values we classified the CD8 T cells into four groups: Cells with a low cytotoxicity score (below the 25^th^ percentile) were classified as naïve or memory-like cells, while the others were considered effector or exhausted if their cytotoxicity scores were significantly higher or lower than expected given their exhaustion scores, respectively. According to this classification, we examined if the clonal expansion program was higher in the effector-like cells. In addition, we compared the SyS CD8 T cells to CD8 T cells from human melanoma tumors (32) using mixed-effects models with a sample-level covariate denoting if the sample was obtained from a SyS or melanoma tumor.

### Malignant epithelial and mesenchymal differentiation programs

The epithelial and mesenchymal signatures were obtained through intra-tumor differential expression analysis, using the likelihood-ratio test for single cell gene expression (69), as implemented in the Seurat package (http://www.satijalab.org/seurat). We compared the mesenchymal to epithelial cells in each of the three biphasic tumor samples (SyS1, SyS12 and SyS12pt). The tumor SyS16 was not included in this analysis (although it was annotated as partially biphasic according to its histology), because its scRNA-Seq sample did not include any epithelial malignant cells. Genes that were up-regulated in the epithelial cells compared to the mesenchymal cells in all three samples were defined as epithelial genes, and likewise for mesenchymal genes. When using the epithelial and mesenchymal signatures in the analysis of bulk gene expression we removed from these signatures those genes that are also part of non-malignant cell type signatures.

Using these signatures we defined: (**1**) the epithelial vs. mesenchymal differentiation score as the OE of the epithelial signature minus the OE of the mesenchymal signature, and (**2**) the differentiation score as the OE of the epithelial signature plus the OE of the mesenchymal signature. An alternative way to define the differentiation score of a particular cell is first to assign it to the epithelial or mesenchymal subset, and then use only the pertaining signature to estimate its differentiation level. However, this approach will not distinguish between poorly-differentiated mesenchymal cells, and mesenchymal cells which have begun to transition to an epithelial state. Hence, we used the inclusive definition of differentiation.

Based on the genes in the epithelial and mesenchymal signatures we then generated diffusion maps (70) for each one of the tumors in our cohort, using the density R package (https://bioconductor.org/packages/release/bioc/html/destiny.html) with the default parameters.

### Identifying co-regulated gene modules

To identify co-regulated gene modules that capture intra-tumor heterogeneity we analyzed each tumor separately. To identify patterns that explain the cell-cell variation both in epithelial and in mesenchymal malignant cells, we further divided the biphasic samples (SyS1, SyS12, and SyS12pt) to their epithelial and mesenchymal compartments. We used PAGODA (71) as implemented in https://github.com/hms-dbmi/scde to filter technical variation and identify co-regulated gene modules in each sample. To identify genes that were repeatedly co-regulated we then constructed a gene-gene co-regulation graph. In this graph, an edge between two genes denotes that the two genes appeared together in the same gene module in at least five samples. Next, we identified dense clusters in the graph using the Newman-Girvan (72) community clustering as previously implemented (73). We filtered out small gene clusters (< 20 genes). Lastly, for each gene cluster we identified the opposing gene module by identifying genes that were negatively correlated with its Overall Expression (OE) across the malignant cells. Correlation was computed using partial Spearman correlation, when controlling for the number of genes and (log-transformed) reads detected per cells, and correcting for multiple hypotheses testing using the Benjamini-Hochberg procedure (74).

For comparison we applied another complementary approach, LIGER (75), which identifies repeating gene modules in the malignant cells using integrative non-negative matrix factorization (NMF) (76). Integrative NMF learns a low-dimensional space, where cells are defined by one set of dataset-specific factors (denoted as *V*_*i*_), and another set of shared factors (denoted as *W*). Each factor, or metagene, represents a distinct pattern of gene co-regulation. To find these metagenes it solves the following optimization problem

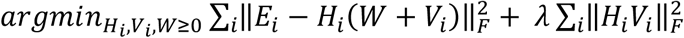

Where *E*_*i*_ denotes the expression matrix (log-transformed TPM) of the malignant cells in sample *i, V*_*i*_ denotes sample-specific metagenes and *W* denotes the shared metagenes across all samples. For this analysis, each biphasic tumor was again split to two “samples”, of epithelial and mesenchymal cells. We used the top 100 genes of each metagene in *W* as the iNMF signatures, and then computed the overall expression of these signatures in the malignant cells. The resulting signatures and their expression across the malignant cells matched the signatures identified with the PCA-based approach, and specifically the core-oncogenic program was re-discovered (**Supp. Fig. 4A**).

### Quantifying RNA velocity

Estimates of RNA velocity were computed using the Velocyto toolkit (http://velocyto.org/). We applied Velocyto with the default parameters, using a gene-relative model. To explore the potential transitions between the epithelial and mesenchymal cell states and avoid confounders, we used only the genes from these differentiation programs (**Supp. Table 4**) for the analysis.

### Predicting patient prognosis

To test if a given program predicts metastasis free-survival or overall survival, we first computed the OE of the program in each tumor based on the bulk gene expression data. Next, we used a Cox regression model with censored data to compute the significance of the association between the expression values and survival. To visualize the predictions of a specific signature in a Kaplan Meier (KM) plot, we stratified the patients into three groups according to the program expression: high or low expression correspond to the top or bottom 20% of the population, respectively, and intermediate otherwise. We used a log-rank test to examine if there was a significant difference between the survival rates of the three patient groups.

### Analysis of *in situ* immunofluorescence imaging

Immune cells were detected based on the protein level of CD45 (>7.5 log-transformed). Malignant cells were identified based on histological morphology, and high protein levels of Hes1. High protein expression was detected by applying the EM algorithm for mixtures of normal distributions. The core oncogenic program score was computed only in the malignant cells based the combined expression of its repressed protein markers: Hsp90, p21, NFkB, and cJun (minus sum of centered log-transformed values). Each image – corresponding to a specific sample in the scRNA-Seq cohort – was divided to frames of 100 cells. The average expression of the core oncogenic program in the malignant cells and the fraction of immune cells in each frame was computed. Using these frame-level values we examined the association between the expression of the core oncogenic program in the malignant cells and the fraction of the immune cells, using a mixed-effects model, with a sample-level intercept (see **Multilevel mixed-effects models**). The mixed-effect model accounts for the nested structure of the data (frames are associated with samples), and ensures the pattern repeatedly appears across different samples.

### Analysis of *in situ* RNA profiling

FASTQ files from multiple lanes were merged to generate single files for processing and insure proper removal of PCR duplicates later in the pipeline. Illumina adapter sequences were trimmed using Trim Galore (version 0.4.5) with a minimum base pair overlap stringency of four bases and a base quality threshold of 20. Paired end reads were stitched using Paired-End reAd mergeR (PEAR, version 0.9.10) specifying a minimum stitched read length of 24bp and a maximum stitched read length of 28bp. The 14bp UMI sequence was extracted from the stitched FASTQ files from the 5’ end of the sequence reads using umi tools (version 0.5.3). The FASTQ files with extracted UMIs were then aligned to a genome containing the 12bp reference sequence tags using bowtie2 (version 2.3.4.1) in end-to-end mode with a seed length of four. Using a custom python function, the generated SAM files were split into multiple SAM files based on the tag to which they aligned to limit memory usage when removing PCR duplicates. The split SAM files were converted to bam files, sorted, and index using samtools (version 1.9) with the import, sort, and index options respectively. PCR duplicates were removed from the sorted and indexed bam files using the dedup command from umi tools with an edit distance threshold of three. An edit distance threshold of three was used. Using custom python functions, the SAM files with PCR duplicates removed were merged for each sample and used to generate digital counts of the tags.

Outlier counts were removed before generating a consensus count for each target. Outlier tags were identified as those with counts 90% below the mean of the probe group in at least 20% of the ROIs analyzed and removed them from the analysis. Subsequently, we removed tags from the analysis if they were flagged as outliers in at least 20% of the AOIs analyzed. This was done using the Rosner Test if there were at least 10 probes for the target (k = 0.2 * Number of Probes, alpha = 0.01), or the Grubbs test if there were less than 10 probes for the target. Probes flagged as outliers in less than 20% of the ROIs analyzed were only removed from the analysis for the ROIs in which they were flagged. Count reported for each target transcript were calculated as the geometric mean of the remaining probes.

The counts for each target transcript were then normalized to the count of the house keeper genes (*C1orf43, GPI, OAZ1, POLR2A, PSMB2, RAB7A, SDHA, SNRPD3, TBC1D10B, TPM4, TUBB, UBB*). The geometric mean of the house keeper gene counts was calculated for each ROI. These geometric means were then divided by the geometric mean of the geometric mean of the house keeper genes to generate a normalization factor for each ROI. The counts of the transcripts in each AOI were than multiplied by the associated normalization factor.

The normalized in situ RNA measures were used to compute: (1) the T cell levels as described in the ***Inferring tumor composition*** section; (2) the overall expression of the malignant programs in each of the regions of interest (ROI), as described in the ***Gene sets Overall Expression*** section; and (3) the differentiation scores, as described in the ***Malignant epithelial and mesenchymal differentiation programs*** section.

### Identifying SS18-SSX targets

The fusion program consists of genes that were differentially expressed in the Aska or SYO1 cells with the SS18-SSX shRNA (shSSX) compared to those with control shRNA (shCt) after 3 or 7 days post-infection. Gene that were previously reported (18,19) to be bound by the SS18-SSX oncoprotein in at least two SyS cell lines were defined as direct SS18-SSX targets, and were used to stratify the SS18-SSX program to direct and indirect targets.

### Mapping cancer-immune interactions

The association between the core oncogenic program in the malignant cells and the expression of different ligands/cytokines in the immune cells was examined using the multilevel mixed-effects regression model described above, using the scRNA-Seq data collected from SyS tumors. The dependent variable *y* was the OE of the core oncogenic program and the covariate *x* was the average expression of a certain ligand/cytokine in a specific type of immune cells (*e.g.*, macrophages) that were profiled from the same tumor. The model also corrected for inter-patient dependencies using the patient-specific intercepts and for cell complexity (log(number of reads)). We restricted the analysis to ligands/cytokines that can physically bind to proteins expressed by the malignant cells (77). The immune cells were either macrophages or CD8 T cells, as other immune cell types were not sufficiently represented in the data.

We used a similar approach to further stratify the program to its TNF/IFN-dependent and independent components. We repeated the same analysis described above, using each one of the genes in the core oncogenic program as the dependent variable. Genes which were associated with both TNF and IFN (P < 0.05, following Bonferroni correction) were considered as TNF/IFN-dependent, and genes which were not associated with both cytokines (P > 0.05) were considered as TNF/IFN-independent.

### TNF and IFN*γ* impact on SyS cell cultures

SyS cell cultures were treated with TNF and IFN*γ*, separately and in combination (see **In vitro IFN/TNF experiment** section), and profiled with scRNA-Seq. Given this data, differentially expressed genes and gene sets were identified using mixed-effects regression models (**Multilevel mixed-effects models** section), with experiment-specific intercepts. The dependent variable *y* was the expression of a gene or the OE of a gene set. The model included three treatment covariates: only TNF, only IFN, and a combination of TNF and IFN. Another binary covariate denoted the duration of the treatment (1 for < 24h duration and 0 otherwise). The model corrected for differences between the different SyS cultures and experiments, and identified patterns that repeatedly appeared across the different experiments. The effect-size and significance of the combination covariate denotes the effect of the combination, and not the synergy between the two cytokines.

To examine if the combined treatment with TNF and IFN*γ* had synergistic effects, we used only the control cells and the cells treated for 4 days with one or two of the cytokines. This model also included 3 binary treatment covariates (TNF, IFN, and the combination), but this time cells that were treated with the combination were positive for all three treatment covariates. The effect-size and significance of the combination covariate hence denotes the synergistic effect of the combination.

### Reconstructing regulatory networks

To reconstruct the gene regulatory network controlling the core oncogenic program we assembled a database of transcription factor (TF) to target mapping based on four sources: JASPAR (78), HTRIdb (79), MSigDB (80,81), and TRRUST (82), and augmented it with the direct SS18-SSX targets identified here (**Supp. Table 5A**) and TF-target pairs we identified in a *cis*-regulatory motif analysis of the core oncogenic program. Specifically, for the *cis*-regulatory analysis, we used RcisTarget (83) (a R/Bioconductor implementation of icisTarget (84) and iRegulon (85)) to identify *cis*-regulatory elements significantly overrepresented in a window of 500bp around the transcription start site of the core oncogenic genes (normalized enrichment score > 3.0) along with their cognate TFs.

We pruned the resulting network to include only core oncogenic program genes (and SS18-SSX) (*i.e.*, all TFs and targets aside from SS18-SSX are program genes). An edge in the network between a TF and its target denotes that: (1) the TF is regulating the target according to at least one of the sources described above, and (2) there is an association between their expression levels in the scRNA-Seq data of SyS tumors. Edges are weighted 1 and −1 to reflect positive and negative associations. We used pageRank (86) (with the R implementation as provided in igraph (https://igraph.org/r/)) as a measure of TF and target importance in the network. To compute TF importance we first flipped the direction of the edges in the network, going from target to TFs. Consistent with the network weights, targets from the up- or down-regulated side of the network were considered induced or repressed, respectively. Likewise, TFs from the up- or down-regulated side of the network were considered activators and repressors, respectively.

### Selectivity and synergy in drug experiments

To evaluate the impact of each drug on the expression of a certain program or gene in a specific cell lines (SYO1, HSSYII, or MSCs), we used a regression model with four binary treatment covariates: abemaciclib, TNF, panobinostat, and the combination of all three drugs. As in the case of TNF/IFN analysis, to examine the synergy of the combination, the cells treated with the combination were positive for all four treatment covariates. The model also included the number of reads detected in each cell (log-transformed) to control for technical variation. When examining the impact on the two SyS cell lines together, we used a mixed-effects model with a cell line specific intercept, to control for cell line specific baseline states. Drug selectivity was examined by using a mixed-effects model that accounts for all three cell lines and has another covariate to denote if the treated cells were SyS or not.

### Data availability

Processed scRNA-seq data and interactive plots generated for this study will be provided through the Single Cell Portal. The processed scRNA-seq data will be provided via the Gene Expression Omnibus (GEO).

## SUPPLEMENTAL FIGURE LEGENDS

**Supplementary Figure 1.**
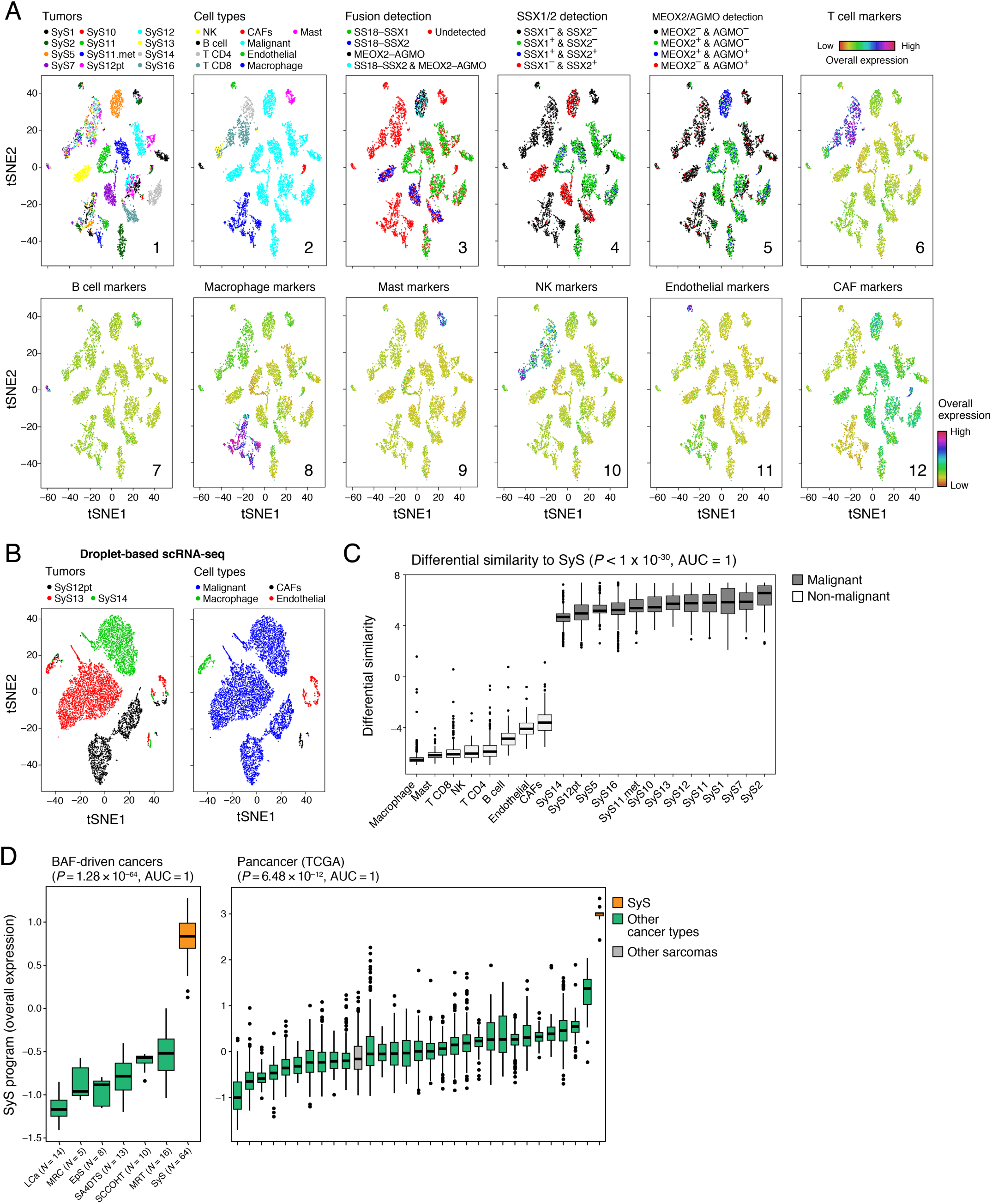
Consistent classification of cells based on expression and genetic features. **(a)** Converging assignments of cell identity. tSNE of single-cell profiles (dots), colored by **(1)** tumor sample, **(2)** inferred cell type, **(3)** SS18-SSX1/2 and MEOX2-AGMO fusion detection, **(4)** *SSX1/2* gene detection (mRNA level > 0), **(5)** *MEOX2* and *AGMO* gene detection (mRNA level > 0), **(6-12)** overall expression of well-established cell type markers (**Supp. Table 2**). **(b)** Droplet based scRNA-Seq of SyS. tSNE of single cells (dots), profiled with droplet-based scRNA-seq (25), colored according to tumor sample (left) and inferred cell type (right). **(c)** Differential similarity to SyS compared to other sarcomas (**Methods**) distinguishes malignant from non-malignant cells. Differential similarity (*y* axis) to SyS shown for cells in each cell subset (*x* axis). Middle line: median; box edges: 25th and 75th percentiles, whiskers: most extreme points that do not exceed ±IQR*1.5; further outliers are marked individually. **(d)** The SyS program distinguishes between SyS and non-SyS cancer types. Distribution of the SyS program Overall Expression (*y* axis) across BAF driven tumors (left, *x* axis) and in TCGA (right, *x* axis). In (c-d) Middle line: median; box edges: 25th and 75th percentiles, whiskers: most extreme points that do not exceed ±IQR*1.5; further outliers are marked individually; P-value: Wilcoxon-ranksum test; AUC: Area Under the receiver operating characteristic Curve.

**Supplementary Figure 2.**
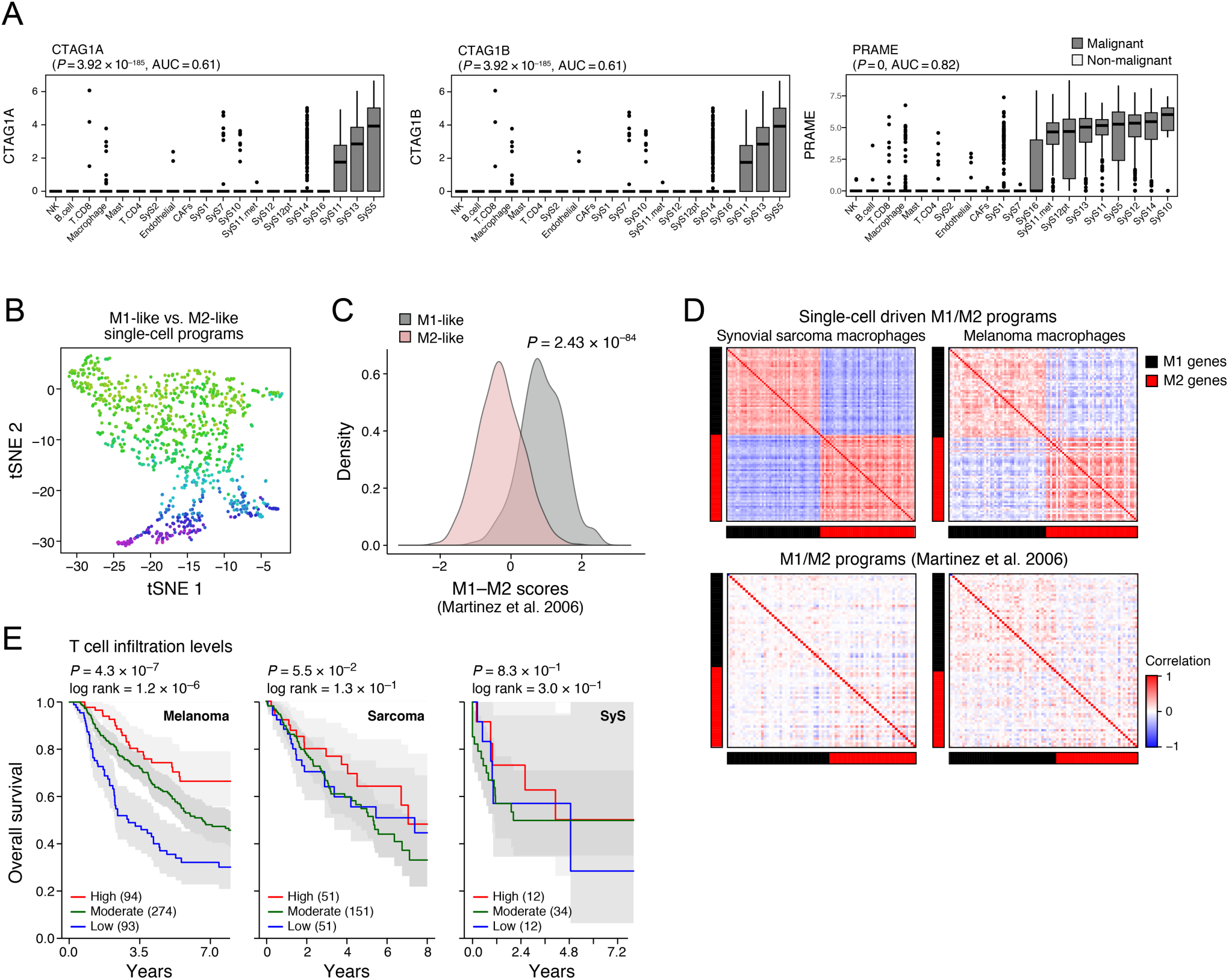
Antitumor immunity and immune evasion in synovial sarcoma. **(a-c)** M1 and M2 states in macrophages are comparable to melanoma. (a) tSNE of macrophage profiles, colored by M1/M2 polarization scores (Overall Expression of the M1 minus M2 program), according to signatures defined here by comparing between the two macrophage clusters (**Supp. Table 3B**). (b) Distribution of M1/M2 polarization scores (*y* axis) according to previously defined signatures (87) in macrophages in our datasets partitioned to M1-like and M2-like subgroups. (c) Spearman correlation coefficient (color bar) between each pair of genes from M1 and M2 signatures defined here (top, **Supp. Table 3B**) or previously (87) (bottom) across macrophages in SyS (left) and melanoma (32) (right). **(d)** Prognostic value of T cell levels in different tumor types. Kaplan-Meier (KM) curves of survival in melanoma (left) (88), sarcoma (middle) (23), and SyS (*8*) (right), stratified by high (top 25%, red), low (bottom 25%, blue), or intermediate (remainder, green) levels of inferred T cell infiltration levels (**Methods**). *P*: COX regression p-value. **(e)** The cancer testis antigens *CTAG1A, CTAG1B* (encoding for NY-ESO-1), and *PRAME* are exclusively expressed by SyS malignant cells. Distribution of expression of each gene (*y* axis, log-transformed TPM) in the cells of each subset (*x* axis). Middle line: median; box edges: 25th and 75th percentiles, whiskers: most extreme points that do not exceed ±IQR*1.5; further outliers are marked individually.

**Supplementary Figure 3.**
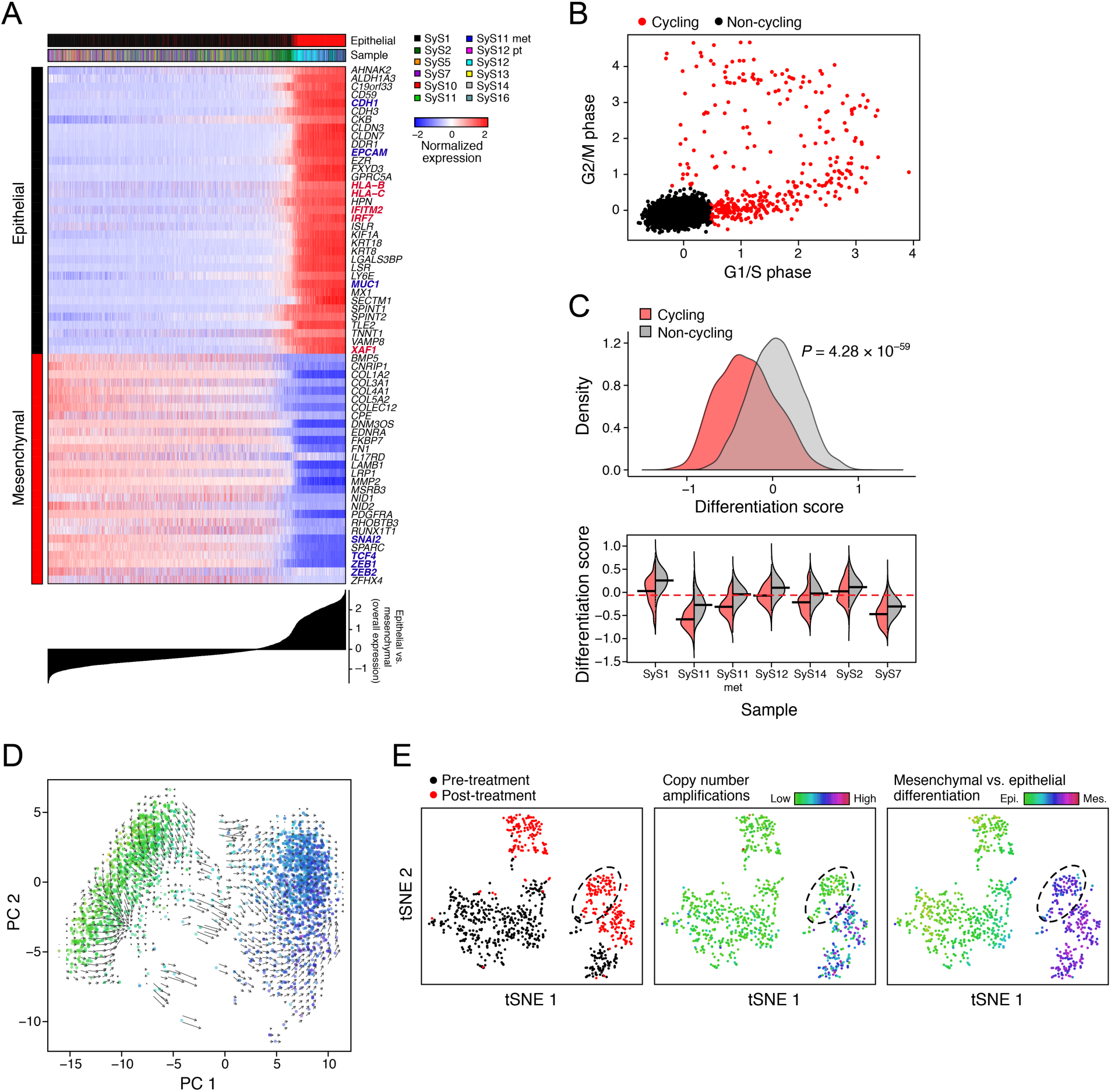
Characterizing mesenchymal, epithelial and poorly differentiates malignant cells. **(a)** Epithelial and mesenchymal program genes. The expression of the top epithelial and mesenchymal program genes (rows) across the malignant cells (columns), with cells sorted according to the difference in epithelial *vs*. mesenchymal OE scores (bottom plot). Topmost Color bar: epithelial *vs*. non-epithelial cell status, and sample. Canonical markers and immune-related genes are in red and blue, respectively. **(b)** Cell cycle signature. Overall Expression of the G2/M (y axis) and G1/S (*x* axis) phase signatures in each malignant cell, colored by their cycling status. **(c)** Cycling cells are less differentiated. The distribution of differentiation scores of cycling (red) and non-cycling (grey) malignant cells, across all tumors (top) and within each tumor (bottom; only tumors with at least 10 cycling cells are shown). **(d)** RNA velocities (38) are visualized on top of the two first principle components (PCs), showing the state and velocity of the malignant cells obtained from patient SyS12 using the droplet-based approach (25). **(e)** t-SNE plots of malignant cells obtained from patient SyS12 before and after treatment, revealing a subpopulation of mesenchymal cells without copy number amplifications in chromosomes 15, 18 and 19 (**Fig. 1G**).

**Supplementary Figure 4.**
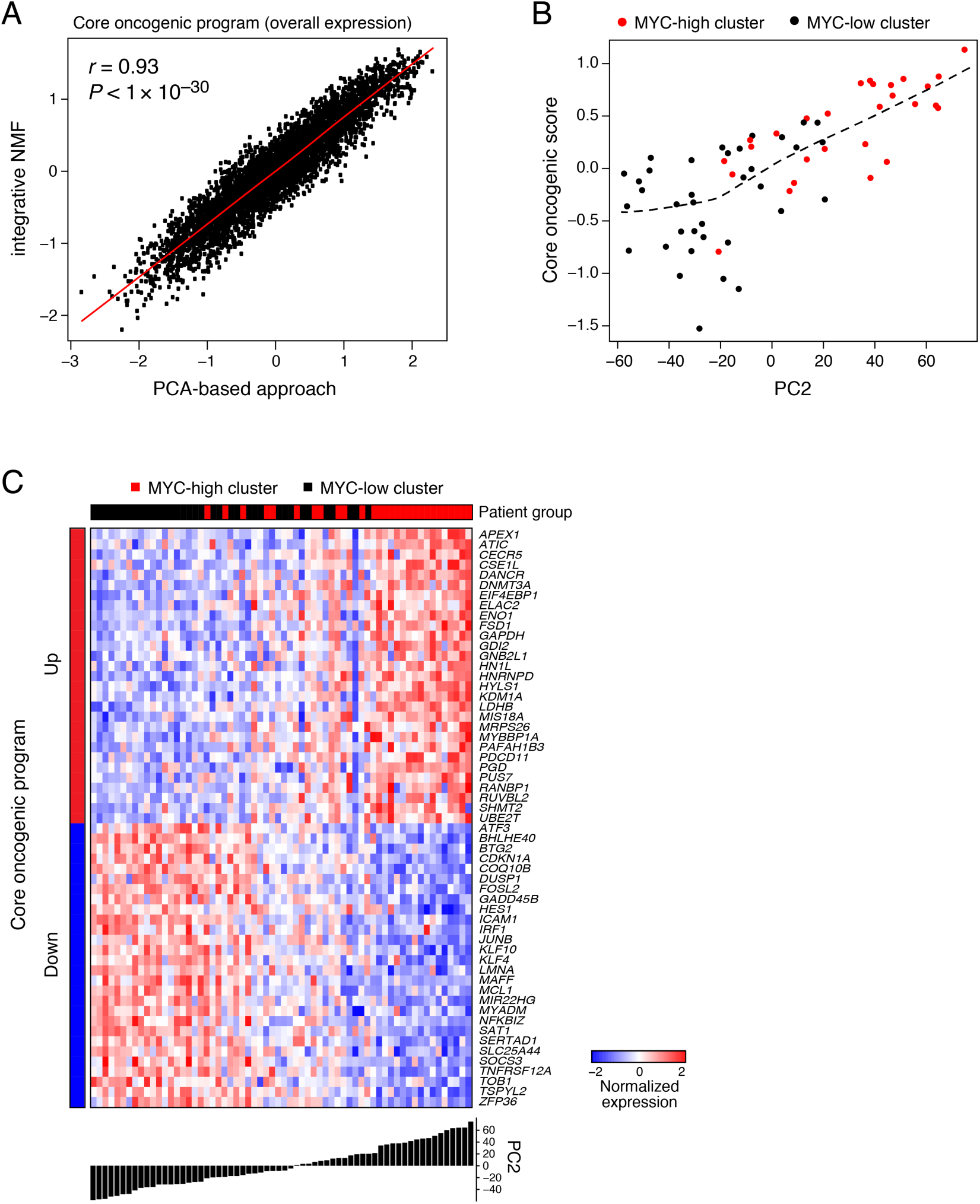
The core oncogenic program is detected using different approaches and datasets. **(a)** Agreement between the core oncogenic program detected by a PCA and an iNMF approach (76). Overall Expression (OE) of the core oncogenic program across malignant SyS cells, as identified in the PCA-based approach (71) (*x* axis) and in the integrative-NMF approach (76) (y axis) (**Methods**). **(b-c)** Program Overall Expression captures inter-tumor variation and the *MYC*-high cluster in 64 SyS tumors from an independent RNA-Seq cohort (18). The tumors were previously classified into two transcriptionally distinct clusters (18), denoted here as *MYC*-high and *MYC*-low. (b) For each tumor (dots), shown is the Overall Expression (OE) of the core oncogenic program (*y* axis) *vs*. the projection on the second Principle Component (PC2) of the data. (c) Normalized expression (centered log-transformed RPKM) of the core oncogenic program genes (columns) most correlated with PC2 across the tumors (columns). Tumors are sorted by their PC2 projection (bottom bar).

**Supplementary Figure 5.**
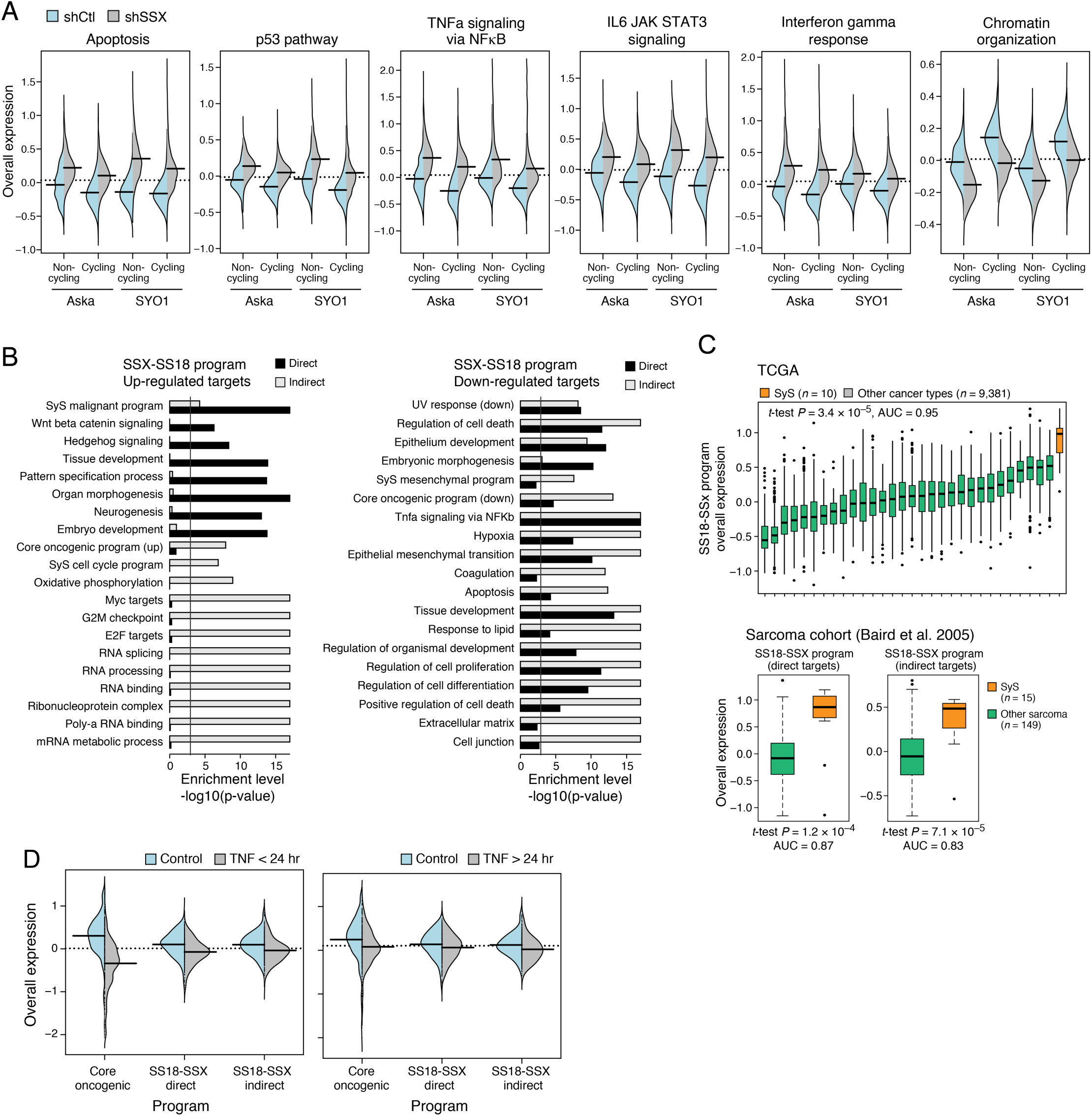
Characterizing the transcriptional impact of SS18-SSX inhibition and tumor microenvironment cytokines on synovial sarcoma cells. **(a)** The fusion KD induces innate immune programs. Distribution of Overall Expression scores (*y* axis) in the pathways most differentially expressed between SyS cells with SS18-SSX (shSSX, grey) *vs*. control (shCt, blue) shRNA, shown separately for non-cycling and cycling cells (*x* axis). **(b)** Biological processes regulated in the SS18-SSX program. Gene sets (rows) most enriched (-log_10_(P-value), hypergeometric test, *x* axis) in induced (left) and repressed (right) SS18-SSX program genes, which are either direct (black bars) or indirect (grey bars) targets of SS18-SSX based on ChIP-Seq data (18,19) and genetic perturbation. Vertical line denotes statistical significance following multiple hypotheses correction. **(c)** The SS18-SSX program distinguishes SyS from other cancer types and other sarcomas. Overall Expression of the SS18-SSX program (y axis) in either TCGA samples (n = 9,391, top), stratified by cancer types (x axis), or in another independent cohort of sarcoma tumors (n = 164, bottom) (48). Middle line: median; box edges: 25th and 75th percentiles, whiskers: most extreme points that do not exceed ±IQR*1.5; further outliers are marked individually. **P<0.01, ***P<1*10^-3^, ****P<1*10^-4^, t-test. **(d)** Repression of the core oncogenic and SS18-SSX programs by short term TNF treatment is not sustained long term. Distribution of Overall Expression scores (*y* axis) of the core oncogenic program and the direct and indirect SS18-SSX programs (*x* axis) in control cells (blue) and cells treated with TNF for 4-6 hours (right) or more than 24 hours (left).

**Supplementary Figure 6.**
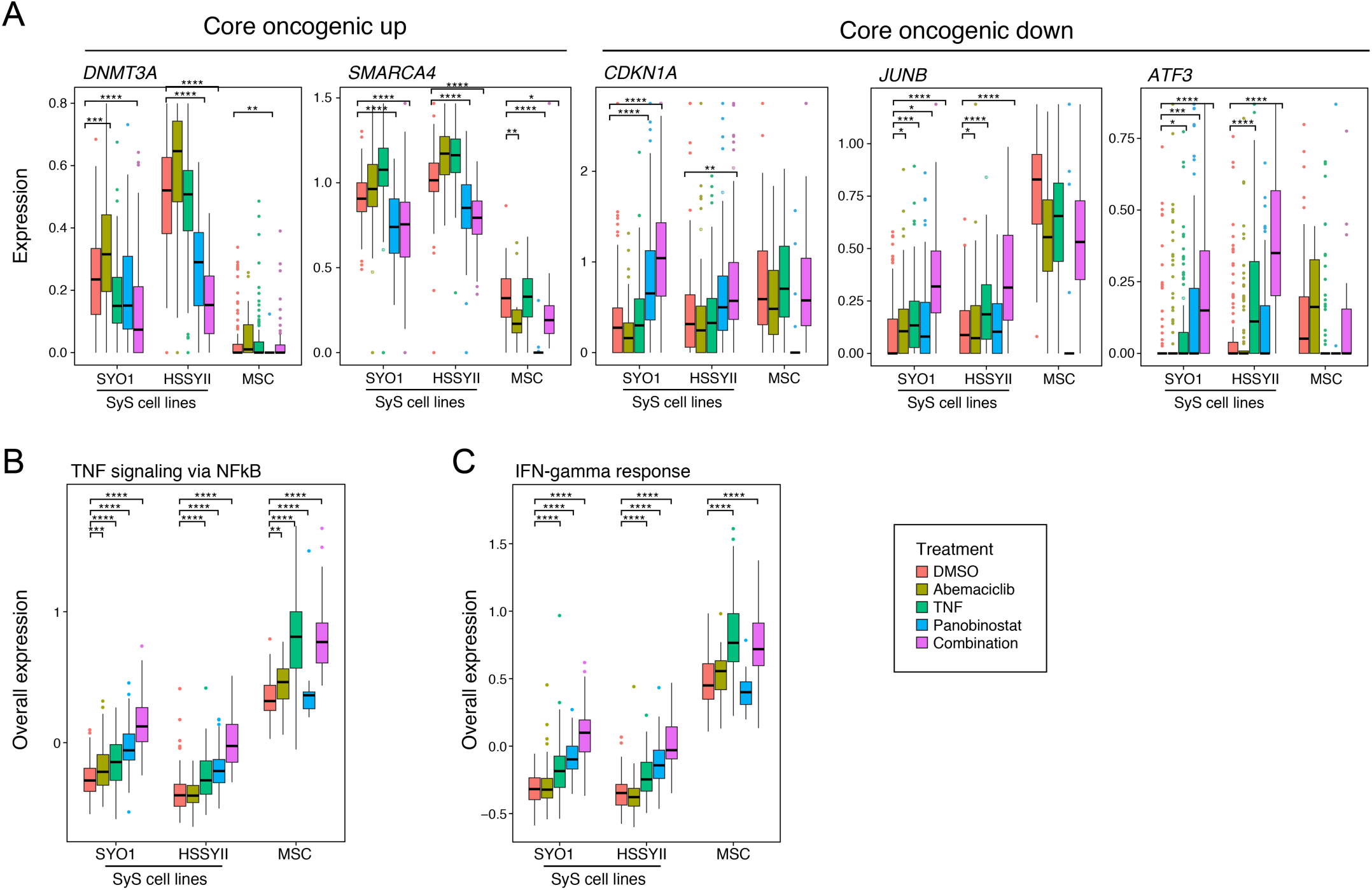
HDAC and CDK4/6 inhibitors synergistically repress the core oncogenic program and induce cell autonomous immune responses. Distribution of the expression (*y* axis) of core oncogenic genes **(a)**, as well as the Overall Expression of TNF **(b)** and IFN **(c)** signaling pathways in SyS cells and MSCs (*x* axis) under different treatments (color legend). Middle line: median; box edges: 25th and 75th percentiles, whiskers: most extreme points that do not exceed ±IQR*1.5; further outliers are marked individually. **P<0.01, ***P<1*10^-3^, ****P<1*10^-4^, t-test.

## SUPPLEMENTAL TABLES LEGENDS

The Supplemental Tables are provided in separate (Excel) files.

**Supplementary Table 1.** (A) Clinical characteristics of the patients and samples in the scRNA-seq cohort and (B) Quality measures of the scRNA-seq cohort.

**Supplementary Table 2.** Cell type signatures derived from the analysis of the SyS scRNA-seq cohort, as well as canonical cell type markers used for cell assignments.

**Supplementary Table 3.** Immune cell states: (A) the T cell expansion program, and (B) M1-like and M2-like macrophage signatures.

**Supplementary Table 4.** Malignant programs: epithelial, mesenchymal, cell cycle and core oncogenic programs (A), and their enrichment with pre-defined gene sets (81) (B).

**Supplementary Table 5.** The fusion program (A) and its enrichment with pre-defined gene sets (81) (B).

**Supplementary Table 6.** TNF and IFN*γ* effects in synovial sarcoma: (A) The predicted TNF/IFN*γ*-dependent and independent components of the core oncogenic program according to the cell-cell interaction analyses (**Methods**); (B) differentially expressed genes following TNF and IFN*γ* treatment, and (C) their enrichment with pre-defined gene sets (81).

